# Corticospinal neuron subpopulations with opposing dynamics converge to shape spinal interneuron computations

**DOI:** 10.64898/2026.10.21.760744

**Authors:** Michelle Sánchez Rivera, Yuxiao Li, Anan Li, Constantinos Eleftheriou, Roman Baravalle, Anand S. Kulkarni, Phillip D. Chapman, Andrés F. Cadena, Francisco J. Alvarez, Jay B. Bikoff, Salvador Dura-Bernal, Xiangning Li, Ian Duguid

## Abstract

Corticospinal neuron (CSN) activation plays a crucial role in orchestrating goal-directed actions, but the functional role of reduced CSN output, a feature conserved across species, remains unknown. By combining population imaging, projection mapping, and modelling in mice performing a Go/NoGo lever-push task, we reveal that distinct corticospinal neuron subpopulations with opposing dynamics (i.e. increased or decreased activity) are selectively associated with skilled movement execution. Both activity profiles are observed throughout CSN somatodendritic compartments and are spatially organized, with increased activity concentrated in upper layer 5B and decreased activity localized to deeper subregions. Notably, single-neuron projection mapping revealed that opposing CSN signals converge onto shared spinal interneurons, while modelling showed that reduced CSN output drives subtractive modulation of spinal interneuron input-output transformations, avoiding output saturation and enabling flexible scaling of output firing rates. These findings redefine corticospinal function by demonstrating that reduced output in a subpopulation of CSNs is critical for shaping spinal cord computations during skilled movement execution.

## Introduction

The corticospinal pathway transforms our intention to move into goal-directed actions. Descending input from corticospinal neurons shape spinal circuit activity necessary for musculoskeletal control and disrupting CSN output results in loss of limb function, impaired reach precision, and hand discoordination^1–9^. While CSN activation has been extensively linked to movement preparation, execution, and parameterization^1,3,4,10–15^, the functional role of CSN suppression – a ubiquitous feature of CSN dynamics across mammalian species^1,15–18^ – remains poorly understood.

In rodents, CSNs innervate multiple targets including the striatum, thalamus, subthalamic nucleus and brainstem, before converging on spinal cord circuits that drive limb kinematics^2,3,19–21^. Projections from discrete subpopulations of CSNs have been shown to target different cell types in supraspinal areas and display only partial overlap in the intermediate and ventral regions of the cervical spinal cord, suggesting projection target divergence. Importantly, each subpopulation broadcasts similar information to downstream supraspinal and spinal cord circuits, implying that motor specificity emerges from the cell types being targeted and differences in synaptic connectivity and connection strengths^15^. This target-dependent specificity, combined with the topographic organization of CSN spinal projections^2,3,15^, creates a functional circuit architecture in which descending cortical output differentially recruits spinal cord circuits to execute a wide range of motor behaviors.

CSN activation coincides with the onset of muscle contraction^15^, consistent with a role in movement initiation, and persists throughout movement execution, reflecting their continuous involvement in online motor control^1,3,15^. However, this activation-based model of CSN dynamics overlooks the significant proportion of neurons that display a reduction in baseline activity during movement (8 – 35% of task-related CSNs), particularly following motor learning^1,15^. This underscores the need to expand this framework to incorporate functional contributions of reduced CSN output^22^. What remains unclear is the functional role of reduced output and whether it is a ubiquitous feature of motor control. The paucity of data regarding the functional relevance of reduced CSN output stems from previous studies focussing on positive changes in CSN somatodendritic calcium dynamics^3,4^ or from using behavioral paradigms that require a single movement, such as pellet reaching or isometric lever pull or press tasks^1,4,15^, which preclude trial-outcome-based comparisons.

To address this, we trained mice on a Go/NoGo lever push task and employed 2-photon CSN somatodendritic calcium imaging with a focus on detecting bidirectional changes in calcium dynamics. By combining cell-type-specific imaging with single neuron axon mapping, retrograde rabies tracing and computational modelling, we establish a revised framework for corticospinal function, revealing that selective suppression of a corticospinal neuron (CSN) subpopulation is essential for sculpting spinal interneuron computations during skilled movement.

## Results

### Movement-based strategies during Go and NoGo trials

To investigate how CSN activity relates to the execution of goal-directed actions, we first developed an auditory discrimination Go/NoGo lever push task for head restrained mice, building on our prior lever task designs^23,24^. This task design allowed us to explore whether decreased CSN activity relates to skilled movement execution, movement suppression, or is a ubiquitous feature of motor control. Following a brief shaping phase (i.e., learning a lever push-reward association, 2 sessions), mice were trained to wait during a fixed inter-trial interval (ITI, 6 seconds) before being presented with either a 6 kHz auditory Go cue or a 13 kHz auditory NoGo cue delivered in a pseudo-randomized sequence. In Go trials, mice required to push a horizontal lever 4 mm forward from the initial starting position. Upon successful execution a servo-controlled lick spout delivered a 5 µl water reward following a 1 s delay, before being reset to its original position. In NoGo trials, mice had to withhold from pushing the lever for 4 s to receive the same reward. Trial outcomes were categorized as Hits, Misses, False Alarms, and Correct Rejections. False Alarms or spontaneous lever pushes during the ITI resulted in a 7.5 s timeout and lever reset before the next trial began (Figure 1A and S1A).

**Figure 1.**
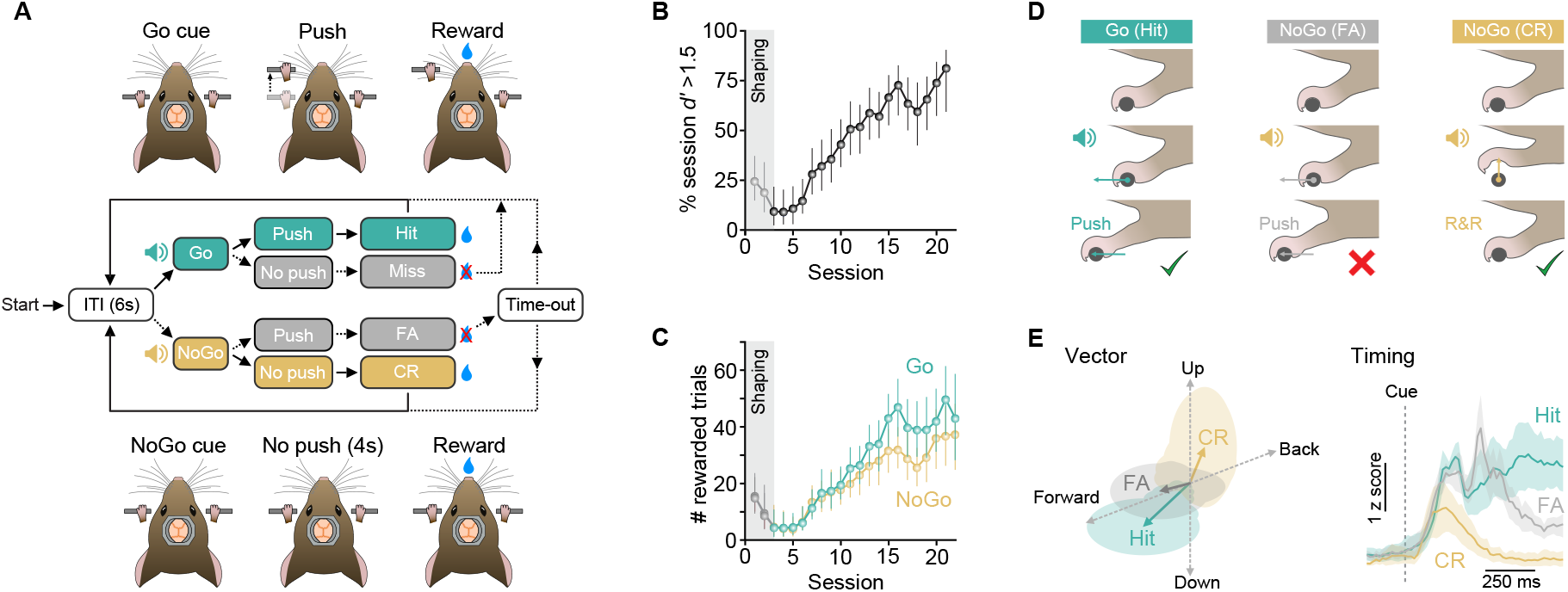
Mice adopt dual movement strategies in a forelimb Go/NoGo lever push task. (A)Behavioral task structure: cued forelimb Go/NoGo lever push task for head restrained mice. Go cue, 6 kHz tone; NoGo cue, 13 kHz tone. ITI, inter-trial interval; FA, false alarm; CR, correct rejection. (B)Percentage of behavioural training session where discriminability index (*d’*) > 1.5. Gray shaded region and symbols depict period of behavioral shaping prior to Go/NoGo training (N = 31 mice). Symbols, mean ± 95% CI. (C)Number of rewarded Go (green) and NoGo (yellow) trials in each behavioral session (40 min). Gray shaded region and symbols depict period of behavioral shaping prior to Go/NoGo training (N = 31 mice). Symbols, mean ± 95% CI. (D)Schematic depiction of paw movements during Go Hit (lever push, green), NoGo False Alarm (FA, lever push, gray) and NoGo Correct Rejection (CR, release and re-grasp, R&R, yellow) trials. Green ticks indicate correct response, red cross indicates an inappropriate response. (E)*Left*, average forelimb movement vectors in a subset of mice (N = 7) during Hit (green arrow), False Alarm (FA, gray arrow) or Correct Rejection (CR, yellow arrow) trials. Arrow length reflects the magnitude of each movement type from cue to median reaction time (i.e., time from cue to lever movement onset) (Hit trial, lever push; False Alarm trial, lever push; Correct Rejection trial, release and re-grasp). Shaded regions, SD across mice. *Right*, average magnitude and timing of forelimb movements in a subset of mice during Hit (push, green), False Alarm (push, gray) and Correct Rejection (release and re-grasp, yellow) trials aligned to cue onset. Data represents mean z-scored motion index ± 95% CI (N = 7 mice).

To evaluate task learning, we computed the discrimination index (*d’*), which quantifies the ability of mice to distinguish between Go and NoGo cues. The effects of satiation on motivation and performance were accounted for by calculating d’ using a rolling window analysis (10 min bin size, 2 min step size)^25^, where mice that maintained a *d’* > 1.5 (i.e. >75% of correct trials) for at least 50% of the session across three consecutive days were considered ‘expert’ (14 [11, 15] 95% CI days to ‘expert’, N = 31 mice). Only epochs where *d’* > 1.5 were included in further analysis. Mice rapidly learned the task, showing a monotonic increase in the proportion of each session with supra-threshold discrimination (Figure 1B) and a corresponding increase in the number of rewarded Go and NoGo trials (Figure 1C and Figure S1B). At expert, mice exhibited highly reproducible forelimb push trajectories during Go trials, with few misses (Figures 1D-1E and Figures S1B-1E). During NoGo trials, mice adopted an unexpected strategy to avoid pushing the lever; they consistently performed a rapid ‘release and re-grasp’ of the lever, followed by a period of immobility lasting 4 s until reward delivery (Figures 1D - 1E and Figure S1C-S1F, Video 1). During False Alarms, mice incorrectly pushed the lever upon NoGo cue presentation, with onset latencies similar to Go push trials (Figures 1D-1E, Figures S1C-S1E and Video 2). However, these movements often resulted in incomplete (“partial”) pushes, suggesting reduced confidence (Figure S1G). To determine whether lever pushes and release and re-grasps were learned during the task, we recorded their incidence in naïve mice during a single session in the absence of any auditory cue. Without shaping, naïve mice did not execute spontaneous lever pushes but frequently repositioned their forelimbs, mirroring release and re-grasp movements observed in expert mice (Figures S1H–J and Video 3). This suggests that mice employ two distinct motor strategies in our task: a learned skilled object manipulation (lever push) and a timed recruitment of a pre-existing posture-related motor program (release and re-grasp).

### CSNs exhibit trial- and movement-type-specific activity

To investigate how CSN activity relates to the execution of these different movements, we combined 2-photon population calcium imaging with an intersectional viral strategy to selectively express GCaMP in C6-projecting corticospinal neurons that control proximal muscles of the forelimb^3^. To do this, we injected rAAV-retro-Cre into the C6 region of the spinal cord (ipsilateral to the moving limb) and AAV-Syn-FLEX-GCaMP6f into the caudal forelimb area (CFA) of motor cortex (contralateral to the moving limb), targeting the region with the highest density of C6-CSNs as identified by retrograde tracing (centered on 1.6 mm ML and 0.2 mm AP to bregma) (Figures 2A and 2B).

**Figure 2.**
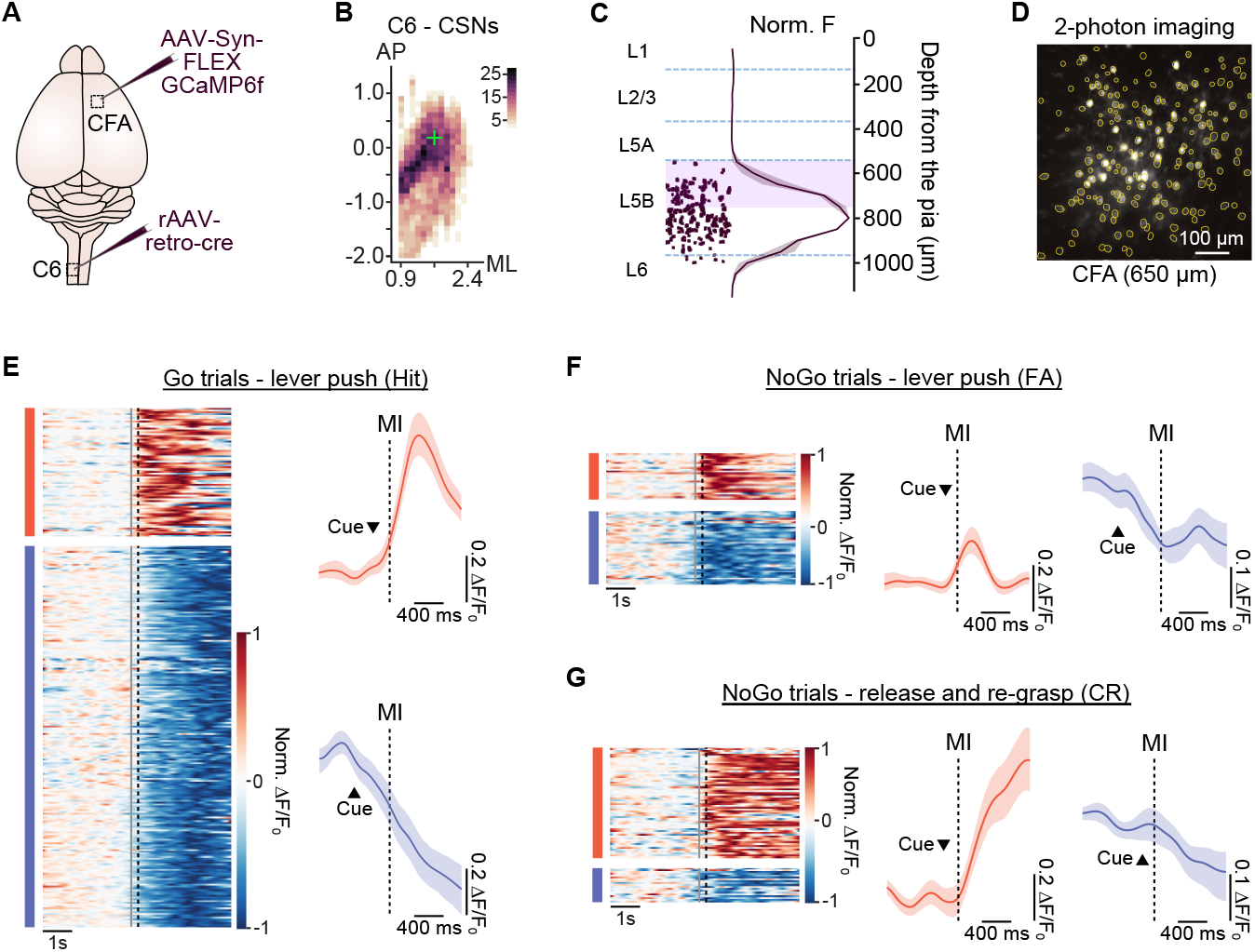
Trial and movement type-specific activity in C6-projecting CSNs. (A)Intersectional viral strategy for expressing GCaMP6f selectively in C6-projecting CSNs. rAAV-retro-Cre injected into the ipsilateral (to the moving paw) C6 segment of the spinal cord and AAV-FLEX GCaMP6f into the contralateral caudal forelimb region (CFA) of motor cortex. (B)Spatial distribution of C6-projecting CSNs in CFA relative to bregma. Scale represents number of cells per bin (bin size = 100 µm AP, 120 µm ML). Green cross, centre of CFA targeted for imaging. ML, medio-lateral; AP antero-posterior. (C)Depth profile of CSNs in CFA. Example slice showing labelling of individual C6-projecting CSNs in layer 5B of CFA and mean normalized fluorescence ± SEM as a function of depth from the pial surface (N = 3 mice). Blue dashed lines depict approximate laminar boundaries; purple shaded area, depth of layer 5B (L5B) accessed by 2-photon imaging. (D)Representative 2-photon imaging field of view (FOV) from layer 5B in CFA. Yellow circles depict regions of interest around identified C6-projecting CSNs. (E)*Left*, Normalized average ΔF/F_0_ for CSNs active during Go trials (lever push) aligned to movement initiation (black dashed line). Each row represents the average activity of an individual CSN. Groupings: increased activity (red bar, n = 71/281 neurons); decreased activity (blue bar, n = 210/281 neurons). In total, n = 281 neurons from 7 fields of view, N = 7 mice. Gray line, cue presentation. *Right*, z-scored ΔF/F_0_ ± 95% CI in example Go trial responsive CSNs showing increased (*top*, red) or decreased (*bottom*, blue) activity when aligned to movement initiation (MI, black dashed line). (F)*Left*, Normalized average ΔF/F_0_ for CSNs active during False Alarm trials (FA, lever push during NoGo trial) aligned to movement initiation (black dashed line). Each row represents the average activity of an individual CSN. Groupings: increased activity (red bar, n = 27/70 neurons); decreased activity (blue bar, n = 43/70 neurons). In total, n = 70 neurons from 7 fields of view, N = 7 mice. Gray line, cue presentation. *Right*, z-scored ΔF/F_0_ ± 95% CI during False Alarm trials in an example CSN showing increased (*left*, red) or decreased (*right*, blue) activity when aligned to movement initiation (MI, black dashed line). (G)*Left*, Normalized average ΔF/F_0_ for CSNs active during NoGo trials (release and re-grasp) aligned to movement initiation (black dashed line). Each row represents the average activity of an individual CSN. Groupings: increased activity (red bar, n = 61/80 neurons); decreased activity (blue bar, n = 19/80 neurons). In total, n = 80 neurons from 7 fields of view, N = 7 mice. Gray line, cue presentation. *Right*, z-scored ΔF/F_0_ ± 95% CI in example NoGo trial responsive CSNs showing increased (*left*, red) or decreased (*right*, blue) activity when aligned to movement initiation (MI, black dashed line).

We monitored CSN population activity at depths between 550 µm and 750 µm from the pial surface, which is amenable to 2-photon imaging (Figures 2C and 2D). To ensure we captured movement-specific activity, we implemented strict inclusion criteria (see Methods) to avoid activity related to reward expectation, reward consumption and post movement postural readjustments (see Figure S1C)^23,24^. Approximately 60% of imaged CSNs (329 / 551 CSNs, 7 fields of view (FOVs), N = 7 mice) exhibited movement-related activity, with around 85% of these showing trial-type specificity (i.e., 281 / 329 CSNs) (Figure S2A – S2D). During lever pushes (Go trial), a minority of CSNs showed increased activity aligned to movement (71 / 281 CSNs, 25%), while the majority displayed decreased activity (210 / 281 CSNs, 75%) (Figure 2E). Notably, CSN activity was movement-type-specific, as no significant ΔF changes were observed during release and regrasps or when data were shuffled (Figure S2A). False alarms, where the lever was inappropriately pushed, recruited a subset of those CSNs active during Go trial pushes. Among these, a larger proportion exhibited decreased activity (43/70, 61%) compared to those with increased activity (27/70, 39%) (Figure 2F). The overlap in CSN responses during Go and False Alarm trials was expected, as both involve lever pushes (Figures 2E and 2F), with the reduced number of active CSNs likely related to the higher probability of partial lever pushes (see Figure S1G). In contrast, during release and re-grasps (NoGo trial), a largely non-overlapping subpopulation of CSNs displayed predominantly increased activity (61/80, 76%), with only a small fraction exhibiting decreased activity (19/80, 24%) (Figure 2G and Figure S2B, S2E). Although previous findings have suggested that baseline ΔF levels predict the direction of movement-related activity changes (i.e. high baseline = decreased CSN activity during movement and vice versa)^1^, we did not observe this relationship. Instead, bidirectional activity changes originated from the same average ΔF/F_0_ baseline (Figures 2E-2G and Figure S2F). Together, these findings indicate that corticospinal neuron subpopulations with opposing dynamics (i.e. increased and decreased activity) are associated with the execution of learned, skilled actions and not a generalized feature of limb movement.

### Temporal and spatial organization of CSN activity

To explore the extent to which increased and decreased CSN activity correlated with limb movement, we first calculated the onset of activity changes using a supervised method for spike inference (CASCADE, see Methods)^26^ and slope sum function (SSF, with parameters adapted to fit our data)^23,27^. During baseline, CSNs exhibited a high level of spontaneous activity, with increased activity tightly time-locked to movement.

Decreased CSN activity was reflected by a rapid reduction in spontaneous events and reduction in mean inferred firing rate that persisted for the duration of movement. Notably, the onsets of increased and decreased activity overlapped prior to movement initiation (−128 [−120, −217] 95% CI ms for increased versus −192 [−160, −256] 95% CI ms for decreased CSNs) (Figures 3A-3D and Figure S3A-S3E), indicative of a synergistic role in initiating skilled movement. In addition, we calculated the bootstrapped median correlation coefficient (Pearson’s r) between the magnitude and timing of forelimb movement (motion index) and deconvolved spike rate (CASCADE) and found that both increased and decreased activity strongly correlated with limb movement during lever pushes (r = 0.74 [0.73 0.74] 95% CI for increased, r = −0.72 [−0.73 −0.71] 95% CI for increased), while increased activity during release and re-grasps was uncorrelated (r = 0.05 [−0.01 0.26] 95% CI) (Figure 3E-3F). This is consistent with CSN activity during release and re-grasps being a cue-driven timing signal related to the execution of a pre-existing posture-related motor program, as evidenced by the lack of activity during spontaneous (or uncued) release and re-grasps performed during the inter-trial interval (Figure S3B).

**Figure 3.**
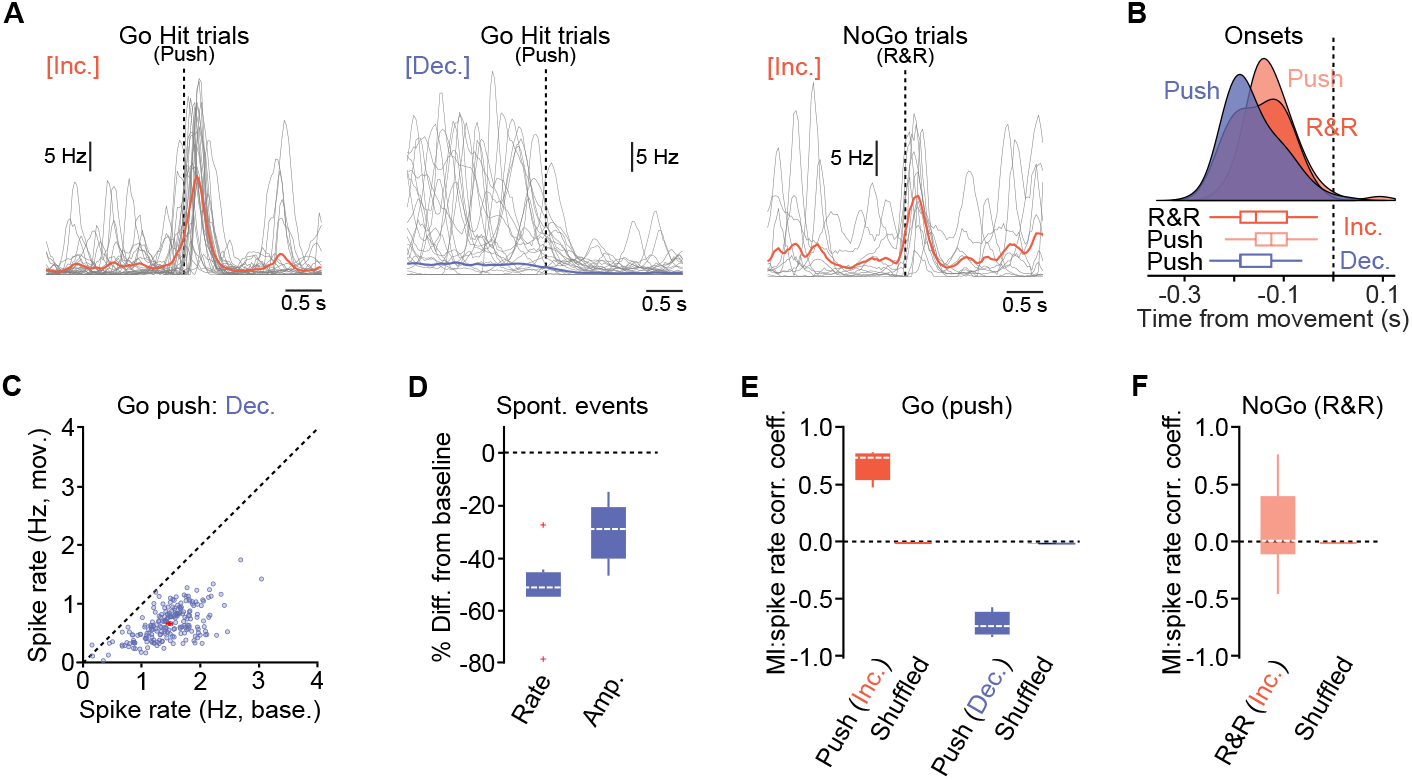
Increased and decreased CSN activity correlates with skilled movement. (A)Overlaid traces of inferred spike rates from CSNs showing (*left*) increased (n = 25 trials overlaid), (*middle*) decreased (n = 21 trials overlaid) activity during Go trial lever pushes or (*right*) increased (n = 11 trials overlaid) activity during NoGo trial release and re-grasps. Dashed line, movement initiation; gray lines, individual trials; thick red or blue lines, mean. (B)Density plot showing onset distributions of firing rate changes during Go trial pushes or NoGo trial release and re-grasps. CSN increased activity, red; decreased activity, blue. Box-and-whisker plots represent bootstrapped estimates of median statistics. Dashed line, movement initiation (N = 6 mice). (C)Inferred spike rates during baseline (base.) and movement (mov.) in CSNs that decreased their activity during Go push trials (n = 210 neurons, N = 7 mice). Red cross, median. Dashed line, unity. (D)Median change in baseline activity in CSNs that show decreased activity during movement. Rate, number of individual bursts of activity. Amp, peak amplitude of individual bursts in Hz (n = 201 neurons, N = 7 mice). Dashed line, median. Red crosses, outliers. (E)Average correlation between forelimb motion index and inferred spike rate in CSNs that increased (red) or decreased (blue) activity during Go trial lever pushes (n = 7 mice). Shuffled data have been added for comparison. Dashed white line, median. (F)Average correlation between forelimb motion index and inferred spike rate in CSNs that increased (red) activity during NoGo trial release and re-grasps (n = 7 mice). Shuffled data have been added for comparison. Dashed white line, median.

Projection neurons in layer 5B of motor cortex receive differential local and long-range inputs depending on depth^28^, which could drive different spatiotemporal activity patterns depending on the type of movement being executed. When observed from a top-down view we found that CSNs displaying increased and decreased activity during lever pushes were spatially intermingled (Figure 4A). However, when viewed along the dorsoventral axis, a clear separation emerged: CSNs with increased activity were predominantly located in the uppermost region of layer 5B, while those with decreased activity were almost exclusively restricted to deeper regions (Figure 4B). This depth-dependent organization was also evident during False alarms, albeit restricted to a small subset of the CSNs active during Go trial pushes (Figures 4C and 4D).

**Figure 4.**
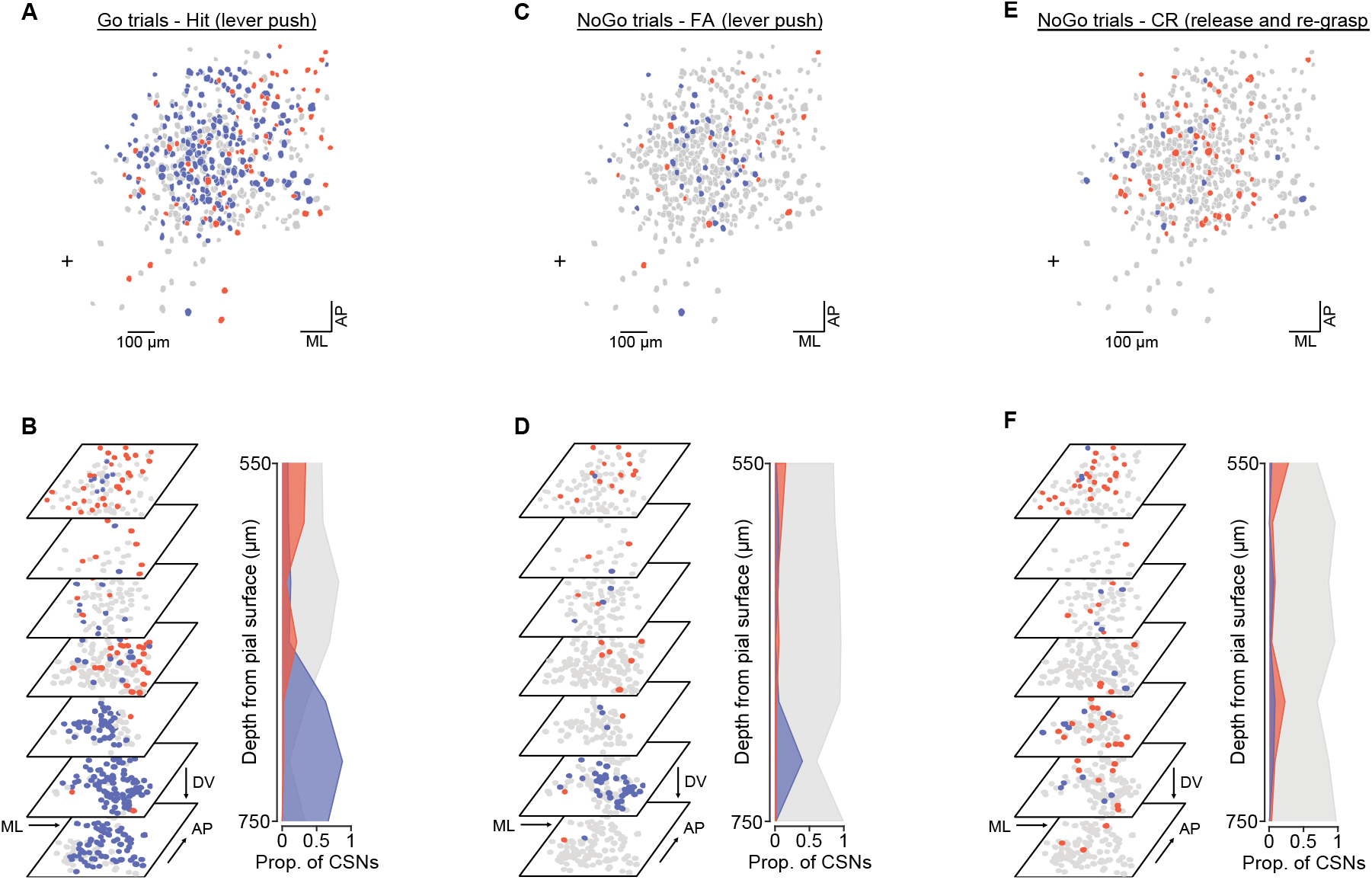
Movement-related CSN activity displays a depth-dependent organization during Go/NoGo task execution. (A)Spatial distribution of CSNs displaying increased (red), decreased (blue) or no change (gray) in activity during Go trial lever pushes (n = 7 overlapping fields of view (FOVs), N = 7 mice). ML, medio-lateral; AP antero-posterior. Black cross, anatomical position from bregma (0 mm AP, 1 mm ML). (B)*Left*, FOVs showing depth-dependent CSN activity patterns (red, increased; blue, decreased; gray no change) during Go trials lever pushes (n = 7 FOVs, N = 7 mice). *Right*, proportion of CSNs displaying increased (red), decreased (blue) or no change in activity (gray) during Go trial lever pushes as a function of depth from the pial surface. ML, medio-lateral; DV, dorso-ventral; AP, antero-posterior. (C)Spatial distribution of CSNs showing increased (red), decreased (blue) or no change (gray) in activity during False Alarm trials (i.e. lever push during NoGo trial) (n = 7 overlapping fields of view (FOVs), N = 7 mice). Black cross, anatomical position from bregma (0 mm AP, 1 mm ML). (D)*Left*, FOVs showing depth-dependent CSN activity patterns (red, increased; blue, decreased; gray no change) during False Alarm trials (i.e. lever push after NoGo cue) (n = 7 FOVs, N = 7 mice). *Right*, proportion of CSNs displaying increased (red), decreased (blue) or no change in activity (gray) during False Alarm trials as a function of depth from the pial surface. (E)Spatial distribution of CSNs showing increased (red), decreased (blue) or no change (gray) in activity during Correct Rejection release and re-grasps (n = 7 overlapping fields of view (FOVs), N = 7 mice). Black cross, anatomical position from bregma (0 mm AP, 1 mm ML). (F)*Left*, FOVs showing depth dependent CSN activity patterns (red, increased; blue, decreased; gray no change) during NoGo trial release and re-grasps (n = 7 FOVs, N = 7 mice). *Right*, proportion of CSNs displaying increased (red), decreased (blue) or no change in activity (gray) during NoGo trial release and re-grasps as a function of depth from the pial surface.

During NoGo trials this dorsoventral gradient of activity was absent with increased activity distributed across both superficial and deeper regions of layer 5B (Figures 4E and 4F). Thus, increased and decreased activity associated with skilled lever pushes in our task are spatiotemporally organized, providing time-locked bidirectional descending input to down-stream targets.

### CSN dendritic activity mirrors movement-related somatic output

We next investigated whether movement specific CSN activity patterns were compartmentalized or widespread across somatodendritic compartments. Pyramidal tract neurons in motor cortex, including CSNs, receive direct somatic input from cortico-cortical and thalamocortical projections, as well as dendritic inputs from local and long-range projections^24,28–30^. Movement-related activity could in principle be driven by direct somatic input without the need for changes in dendritic excitability. Conversely, such changes could be driven by neuron-wide inputs, and thus be visible across all somato-dendritic compartments. To distinguish between these two possibilities, we performed near-simultaneous multi-plane imaging of the apical tuft and proximal dendrites of CSNs during task execution. We employed an intersectional labeling strategy, where diluted rAAV-retro-Cre (1:150) was injected into the C6 spinal cord to achieve sparse labelling, and a Cre-dependent bicistronic vector expressing GCaMP6s and mRuby2 into motor cortex, to record activity and enable full morphological reconstructions^31,32^. CSNs were identified as type I thick-tufted layer 5 pyramidal tract neurons, characterized by an early-bifurcating apical tuft and long nexus (n = 16 neurons, N = 9 mice) (Figure S4B and S4C)^31^. We recorded activity from 10 - 30 dendritic branches per CSN, 22 CSNs, 8 mice, with imaging planes spaced ~100 – 200 µm apart and sampled at 8.33 Hz per FOV. This allowed near-simultaneous recording of multiple dendritic regions of interest (ROIs) within the same CSN over 2 consecutive days (Figures 5A and 5B). ROIs were aligned to the reconstructed dendrites, and activity classifications were based on the average ΔF/F_0_ across branches during different trial types.

**Figure 5.**
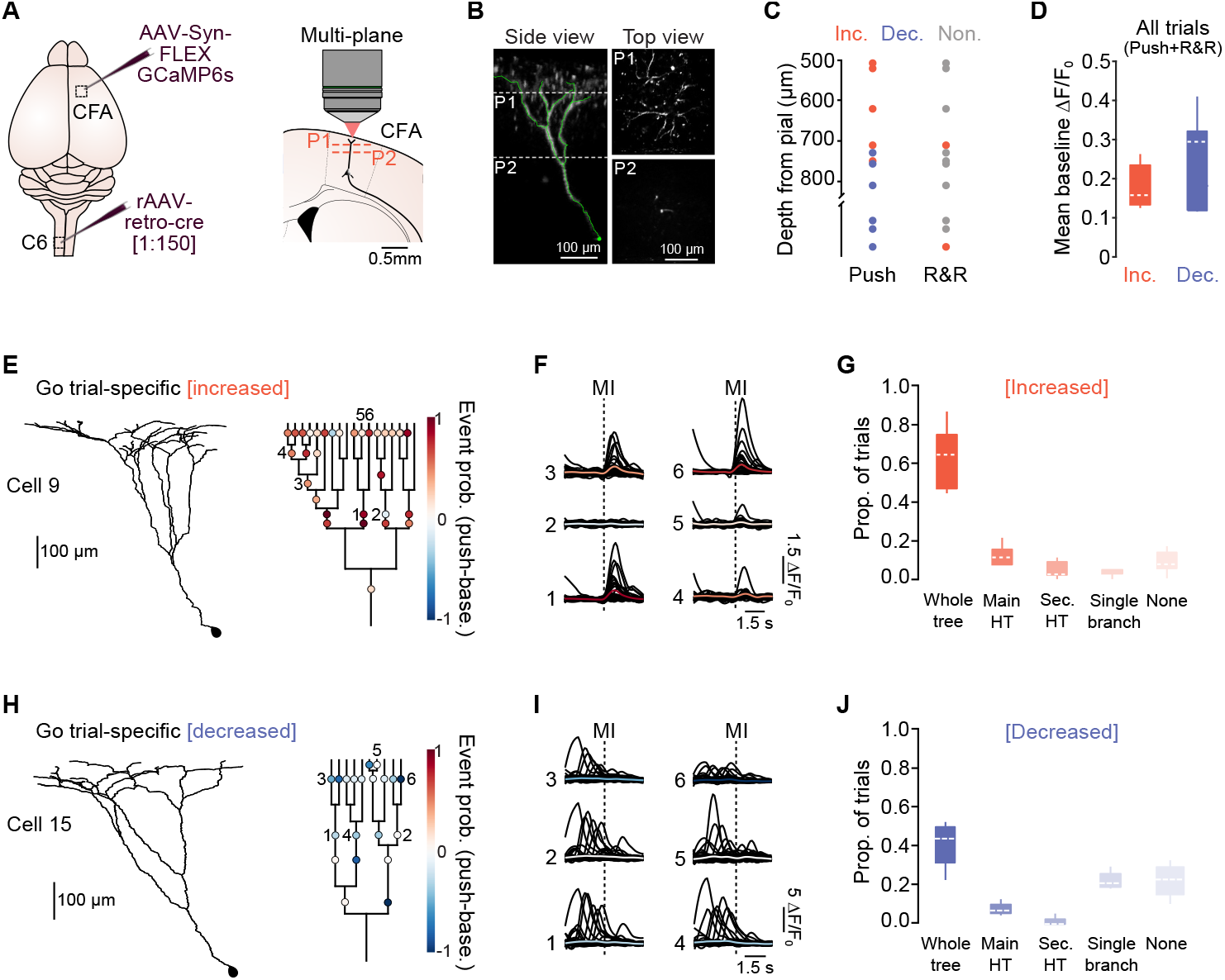
CSN dendritic activity reflects bidirectional trial-type-specific somatic output. (A)*Left*, intersectional viral strategy for expressing GCaMP6s selectively in C6-projecting CSNs. Diluted rAAV-retro-cre [1:150] injected into the ipsilateral (to the moving paw) C6 segment of the spinal cord and AAV-FLEX GCaMP6s into contralateral CFA. *Right*, schematic illustrating multi-plane 2-photon dendritic imaging of C6-projecting CSN dendrites (black). CFA, caudal forelimb region of motor cortex; P1 - first imaging plane; P2, second imaging plane (~ 200 µm between planes, 3-4 planes per CSN). (B)*Left*, 2-photon z-stack image (side view) of a C6-projecting CSN from soma to apical tuft. *Right*, FOVs (top view) at two imaging planes showing apical tuft (P1) and parent dendrites (P2). (C)Somatic depth of C6-projecting CSNs identified as showing increased (red) or decreased (blue) dendritic activity during Go push trials or NoGo release and re-grasps. Gray symbols, no change. (D)Box plots showing the distributions of mean baseline ΔF/F_0_ in CSN dendrites displaying increased (red) or decreased (blue) activity during movement. Note Inc. and Dec. include both Go and NoGo specific CSNs. Dashed line, median. (E)*Left*, reconstruction of the trunk and apical dendrites of a Go trial-specific CSN displaying increased activity during lever pushes. *Right*, corresponding dendrogram showing location of imaged dendrites and change in event probability across trials (i.e. event probability during push - event probability during baseline period). (F)Overlaid raw ΔF/F_0_ traces (black) and mean (colored) from individual dendrites numbered in panel (E) (35 consecutive trials). (G)CSNs with Go trial-specific increased dendritic activity: box plots showing proportion of trials with increased activity during push movements. Activation patterns are separated into whole dendritic tree (whole tree), main hemi-tree (Main HT), secondary hemi-tree (Sec. HT), single branch and trials with no increased activity (None) (n = 5 neurons, N = 4 mice). Dashed line, median. (H)*Left*, reconstruction of the trunk and apical dendrites of a Go trial-specific CSN displaying decreased activity during lever pushes. *Right*, corresponding dendrogram showing location of imaged dendrites and change in event probability across trials (i.e. event probability during push - event probability during baseline period). (I)Overlaid raw ΔF/F_0_ traces (black) and mean (colored) from individual dendrites numbered in panel (H) (47 consecutive trials). (J)CSNs with Go trial-specific decreased dendritic activity: box plots showing proportion of trials with decreased activity during push movements. Activation patterns are separated into whole dendritic tree (whole tree), main hemi-tree (Main HT), secondary hemi-tree (Sec. HT), single branch and trials with no decreased activity (None) (n = 4 neurons, N = 3 mice). Dashed line, median.

The proportions of CSNs exhibiting trial- and movement-type specific dendritic activity closely mirrored somatic activity patterns (11/22 task-active CSNs; 9/11 displayed movement-type specific activity), suggesting a strong correlation between calcium events in the soma and apical dendrites^1,31,33,34^. During lever pushes, dendritic tufts exhibited increased or decreased activity and the mapping of their somatic location again revealed a dorsoventral gradient: CSNs showing increased activity were predominantly located in the uppermost region of layer 5B, while those with decreased activity were restricted to deeper regions. In contrast, release and re-grasps were associated with increased activity in a small number of CSNs with somata distributed across both upper and deeper layer 5B (Figure 5C). Baseline dendritic fluorescence differed between the two CSN subpopulations, with CSNs that reduced their activity displaying slightly higher baseline fluorescence (Figure 5D)1. However, this difference was not reflected at the somatic level (see Figure S2).

To quantify the extent of dendritic activation during movement, we identified four possible activation patterns: whole tree; hemi-tree; single branch; and none (or no branches) (Figure S5A)^31^. In CSNs with increased activity during movement (lever push or release and re-grasp) the predominant spatiotemporal profile was whole tree activation (64% [53% 74%] trials), which involved almost all tuft dendrites (~70% of tuft dendrites per trial). However, a gradient of activation was observed, with dendrites just after the nexus displaying more consistent trial-to-trial activity compared to distal dendrites (Figure 5E). Dendritic activation mirrored the temporal dynamics of somatic activity, rising at movement initiation, and peaking around movement completion (Figure 5F and Figure S5E). Notably, not all trials involved whole tree activation; in ~20% of trials, only one hemi-tree was active, with occasional switching between trials (left to right & vice versa) and in a minority of trials (~5%), activity was restricted to a single dendritic branch (Figure 5E-5G and Figures S4D-4E and S4H). Parallel to dendritic activation, we also observed a large proportion of CSN dendritic tufts that exhibited a rapid reduction in spontaneous activity around movement initiation, with no movement-related activation (44% [40% 48%] silent trials). Reduced activity was observed across the whole dendritic tree with low trial-to-trial variability and persisted for the duration of the movement (Figures 5H-5J and Figures S4F-S4G and S4I). This dampening of spontaneous activity unmasked a higher incidence of single branch events (~25% of trials), indicative of highly localized, branch-specific spiking. Thus, while CSNs dendrites do display compartment-specific activity^31,35,36^, our data suggest a high degree of CSNs somatodendritic coupling whereby movement-related activity is dominated by global, multi-compartmental modulation that strongly correlates with bidirectional somatic output^31,33,34,37^.

### CSN outputs converge on V1 spinal interneurons

Given these findings, we focused the rest of our investigation on understanding how subpopulations of CSN with opposing dynamics (increased vs decreased) might shape downstream spinal cord computations during skilled action execution. Although the observed dorsoventral gradient of activity observed in our task is likely movement specific, we took advantage of this functional anatomical arrangement to explore how this information propagates to the spinal cord. The two subpopulations of CSNs could in principle generate two parallel, non-overlapping streams of information targeting distinct supraspinal and spinal regions (segregation model). Alternatively, they might exhibit substantial overlap and downstream convergence (convergence model) (Figure 6A). To resolve this, we adopted a dual approach: fluorescence micro-optical sectioning tomography (fMOST) to map CSN axonal projections to supraspinal and spinal targets^38,39^, and transsynaptic retrograde rabies virus (RV) tracing from defined cervical spinal interneuron subpopulations, combined with serial two-photon tomography (STPT) to reconstruct the spatial distributions of CSNs^40^.

**Figure 6.**
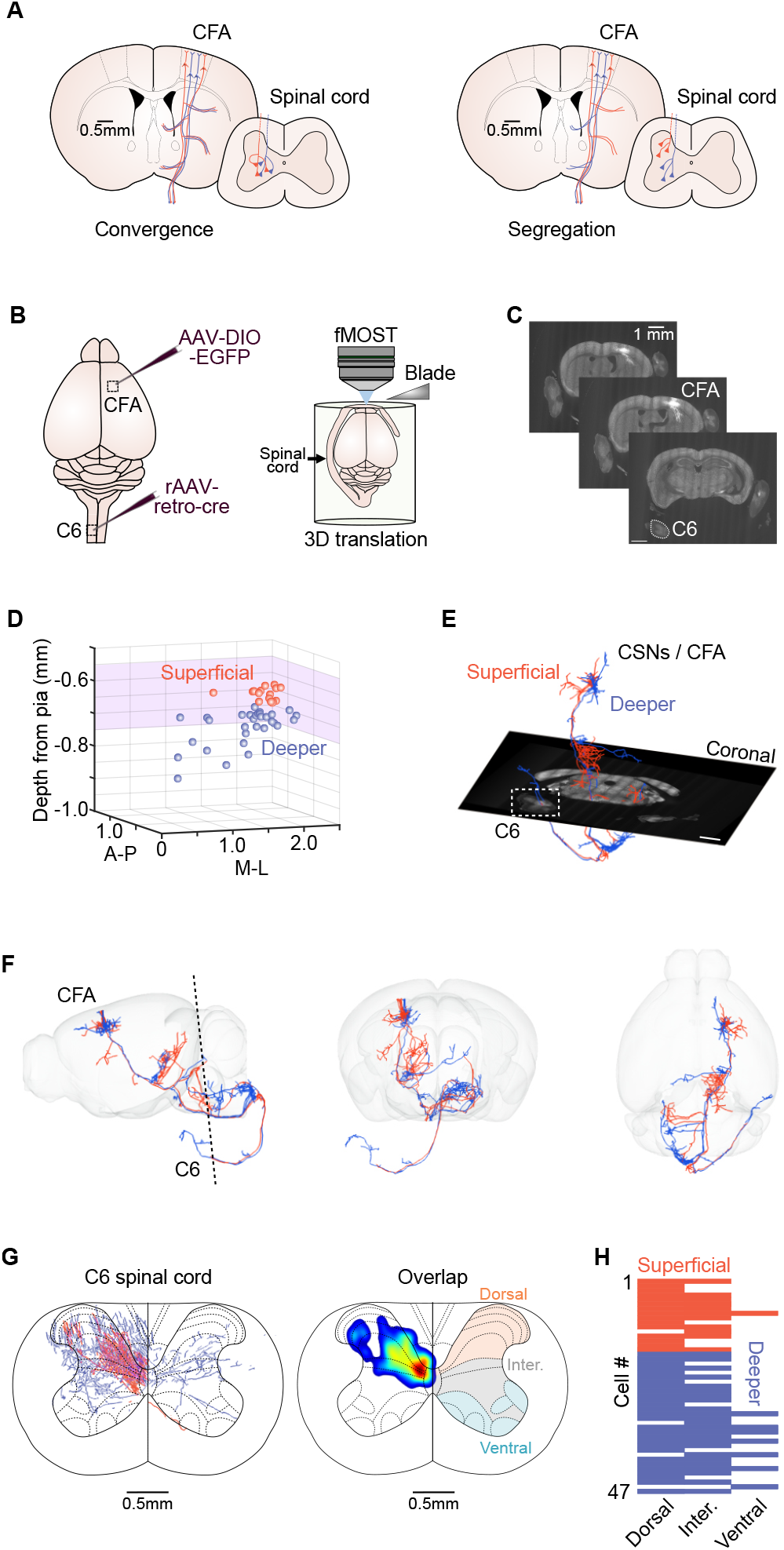
CSN projections from superficial and deeper regions of CFA converge in cervical spinal cord. (A)Schematic diagrams showing supraspinal and corresponding spinal cord projection targets of CSNs in superficial (red) and deeper (blue) regions of layer 5B caudal forelimb area (CFA). *Left*, Convergence model, where projections from CSNs in superficial and deeper regions of CFA target the same supraspinal and spinal cord areas. *Right*, Segregation model, where projections from CSNs in superficial and deeper regions of CFA target different supraspinal and spinal cord regions. (B)fMOST imaging approach. *Left*, intersectional viral strategy for expressing EGFP selectively in C6-projecting CSNs. rAAV-retro-Cre injected into the ipsilateral C6 segment of the spinal cord and AAV-DIO-EGFP into the contralateral caudal forelimb region (CFA) of motor cortex. *Right*, Schematic showing brain and spinal cord preparation embedded in resin where 2 µm slices are cut using a diamond blade and imaged using 1-photon patterned light stimulation. fMOST, fluorescence micro-optical sectioning tomography. (C)Example coronal images of the brain and spinal cord acquired using fMOST Scale bar = 1 mm. (D)Somatic location of reconstructed CSNs in CFA layer 5B. Red, superficial CSNs (550 - 650µm); Blue, deeper CSNs (650-900µm). Purple shaded area, region imaged using 2-photon microscopy as shown in Figure 2. ML, medio-lateral; AP antero-posterior. (E)Example reconstructions of CSNs located in superficial (red) and deeper (blue) regions of CFA superimposed on a coronal brain / spinal cord slice. White box depicts axon reconstructions in the C6 region of the spinal cord. Scale bar = 1 mm. C6, C6 segment of the spinal cord; CFA, caudal forelimb region of motor cortex. (F)Example visualizations of two CSNs from superficial (red) and deeper (blue) regions of CFA. Dashed line, approximate imaging plane shown in (E). CFA, caudal forelimb region of motor cortex. (G)*Left*, spatial distribution of CSN axons in the C6 region of the spinal cord (n = 47 CSNs, N = 4 mice) where red axons are from superficial CSNs and blue axons from CSNs deeper in layer 5B CFA. *Right*, axon density overlap in intermediate (inter.) and dorsal regions of spinal cord. Note, highest density overlap in intermediate zone. (H)Raster plot showing proportion of CSNs in superficial (red) and deeper (blue) regions of CFA that target dorsal, intermediate and ventral regions of the spinal cord (n = 47 CSNs, N = 4 mice). Note, CSNs are ordered by depth from the pial surface.

First, we labelled C6-projecting CSNs (n = 47 neurons, N = 4 mice) using our intersectional viral strategy with AAV-DIO-EGFP injected into CFA to facilitate morphological reconstruction of CSN axons. Whole brain and spinal cord preparations were embedded in resin and mounted on a 3D translational stage where 2 µm slices were cut using a diamond blade and imaged using 1-photon structured illumination using fMOST.

CSN axons were reconstructed using GTree (see Methods) and aligned to the Allen Mouse Common Coordinate Framework (CCFv3) and SpinalJ template (https://zenodo.org/records/5177185) for visualization (Figure 6B and 6C). Importantly, our sample of 47 fully reconstructed CSNs spanned both superficial (<650 µm) and deeper regions (>650 µm) of layer 5B, consistent with the depth profile classification of our somatic and dendritic recordings (Figure 6D). Supraspinal projections from both CSN subpopulations were largely overlapping in the midbrain, basal ganglia, hypothalamus, pons, and thalamus. However, superficial CSNs exhibited sparse or no axon labelling in the medulla, consistent with the notion that pyramidal tract neurons in deeper layer 5B project primarily to the medulla^41^ (Figure 6E and 6F, Figure S5). In the spinal cord, axons from both superficial and deeper CSNs displayed the highest degree of overlap in the dorsal and intermediate zones of C6 (# of CSNs targeting dorsal region: superficial 15/16, deeper 28/31; # of CSNs targeting intermediate region: superficial 11/16, deeper 24/31, N = 4 mice), with a small proportion of deeper CSNs projecting to the ventral region (8/31 CSNs, N = 4 mice) (Figure 6G and 6H).

CSNs in the caudal forelimb area of motor cortex project to multiple excitatory and inhibitory interneurons in the spinal cord but display a substantial bias towards inhibitory populations, and a specific preference for V1 interneurons found in the intermediate and ventral zones^42,43^. We leveraged this fact and that axon convergence was highest in the intermediate zone to analyze a previously acquired transsynaptic retrograde rabies tracing dataset that mapped the spatial distributions of CSNs providing monosynaptic input to V1 interneurons in the cervical spinal cord (see Figure S3C in ^44^). We reasoned that if both subpopulations of CSNs converge onto V1 interneurons, retrogradely labelled CSNs should span the entirety of motor cortex layer 5B (superficial *and* deep) (Figure 7A). Conversely, if they target separate interneuron populations, labelling would be restricted to either the superficial *or* deeper region CSNs (Figure 7B).

**Figure 7.**
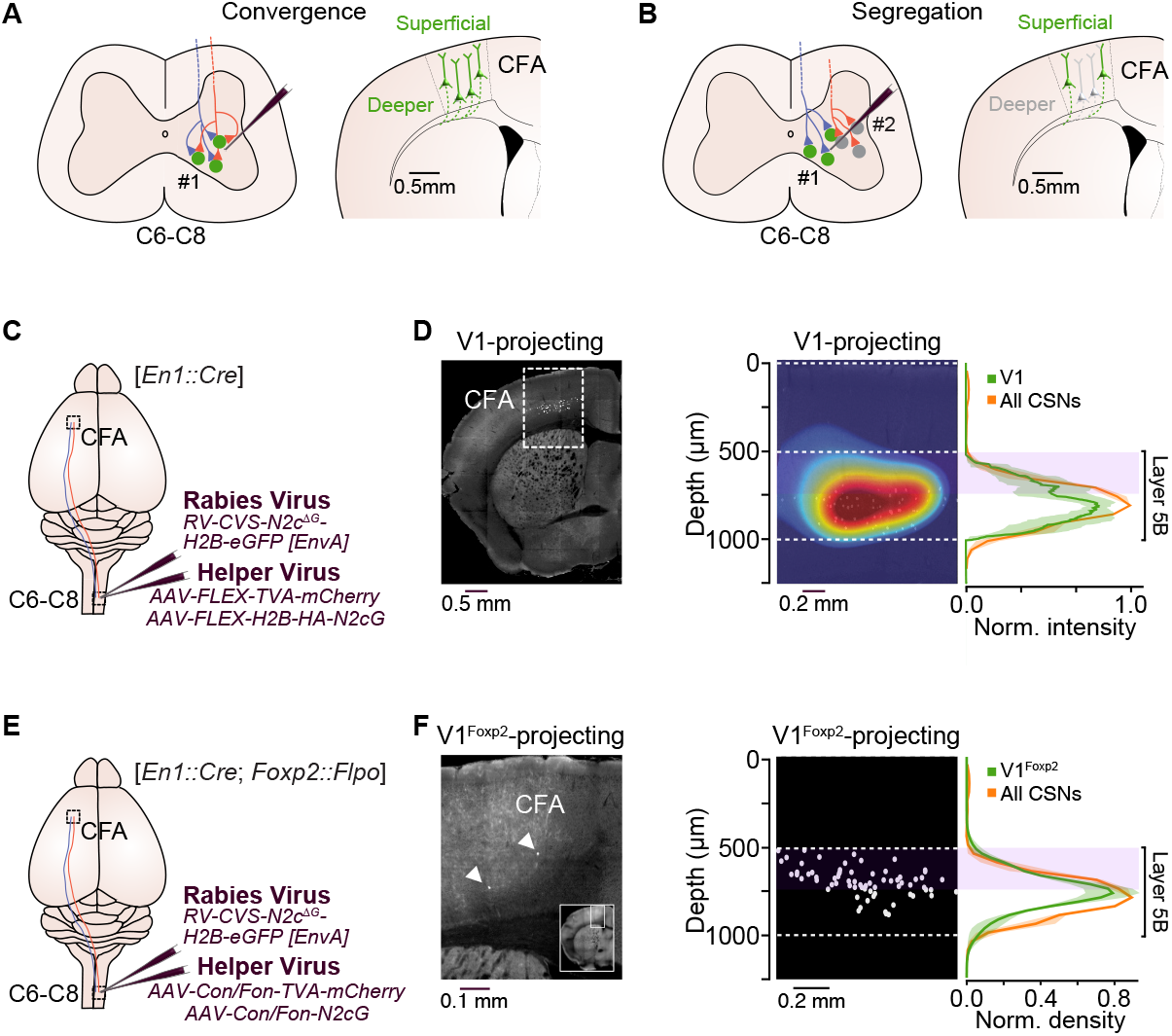
CSN projections from superficial and deeper regions of CFA target V1 spinal interneurons. (A)*Left*, Convergence model: CSN projections from superficial (red) and deeper (blue) regions of CFA target the same spinal interneuron subpopulation. *Right*, using the spinal interneuron subpopulation #1 as starter cells, retrograde rabies tracing labels CSNs in both superficial and deeper regions (green) in CFA. (B)*Left*, Segregation model: CSN projections from superficial (red) and deeper (blue) regions of CFA target different spinal interneuron subpopulations. *Right*, using the spinal interneuron subpopulation #1 as starter cells, retrograde rabies tracing only labels superficial CSNs (green) in CFA. (C)Strategy to label V1 spinal interneuron-projecting CSNs in the C6-C8 region of the spinal cord using transsynaptic rabies tracing. *En1::Cre*, mouse line expressing Cre in V1 spinal interneurons; CFA, caudal forelimb region. Schematic adapted from Chapman et al., 2025. (D)*Left*, retrogradely labelled V1 spinal interneuron-projecting CSNs in CFA. *Right*, heat and density plots showing the location of V1-projecting CSNs (green) and overlap with all retrogradely labelled C6-C8-projecting CSNs (orange) in layer 5B of CFA (N = 3 mice). Dashed lines, upper and lower borders of layer 5B; light purple shading, depth of 2-photon imaging. (E)Strategy to label V1^Foxp2^ spinal interneuron-projecting CSNs in the C6-C8 region of the spinal cord using transsynaptic rabies tracing. *En1::Cre*; *Foxp2::Flpo* mouse line expressing Cre and Flpo in Foxp2 expressing spinal interneurons; CFA, caudal forelimb region. Schematic adapted from Chapman et al., 2025. (F)*Left*, retrogradely labelled V1^Foxp2^ spinal interneuron-projecting CSNs in CFA. White arrows highlight 2 labelled CSNs. *Right*, locations of V1^Foxp2^ spinal interneuron-projecting CSNs (green) and comparison with all retrogradely labelled C6-C8-projecting CSNs (orange) in layer 5B of CFA (N = 3 mice). Dashed lines, upper and lower borders of layer 5B; light purple shading, depth of 2-photon imaging.

The approach involved selectively targeting V1 spinal interneurons by injecting Cre-dependent helper viruses (AAV-FLEX-TVA-mCherry and AAV-FLEX-H2B-HA-N2cG) unilaterally into C6-C8 spinal segments of *En1::Cre* mice, followed 5 weeks later by injection of EnvA-pseudotyped RV (CVS-N2cΔG) expressing histone-tagged H2B-eGFP (Figure 7C). After 1 week to permit monosynaptic retrograde transfer to the brain, labeled CSNs were imaged via STPT and analyzed to determine their precise laminar location. This approach revealed V1-projecting CSNs distributed across the entire dorsoventral axis of CFA layer 5B, matching the distribution of all C6-projecting CSNs (Figure 7D). Notably, the V1 interneuron population is comprised of several different subtypes defined by transcription factor expression, with the Foxp2 subpopulation (V1^Foxp2^) being the preferred target of CSNs from CFA^42^. To explore the laminar distribution of CSNs innervating V1^Foxp2^ interneurons, we performed RV tracing in *En1::Cre*; *Foxp2::Flpo* mice, taking advantage of an intersectional viral strategy that drives TVA-mCherry and N2cG expression through the combined activity of Cre and Flp recombinases (AAV-Con/Fon-TVA-mCherry and AAV-Con/ Fon-N2cG respectively) (Figure 7E)^44^. This approach revealed that V1^Foxp2^-targetting CSNs were also distributed across the entire dorsoventral axis of CFA layer 5B (Figure 7F), providing further support for a convergence model in which both increased and decreased CSN activity converge on V1 interneuron populations in the intermediate zone of the spinal cord.

### Reduced CSN input drives subtractive modulation of V1 input-output transformations

Given this anatomical organization, we sought to understand how bidirectional CSN output might affect activity in the spinal cord using V1 interneurons as an exemplar. Considering the challenges in performing intracellular recordings in the spinal cord of awake mice and difficulty in selectively manipulating either increased or decreased CSN activity, we instead opted for a computational approach by developing a multi-compartment biophysical model of a V1 interneuron based on *in vitro* slice recordings (see Methods)^45^. Our model recapitulated the current-frequency relationship, moderate peak firing rates and tendency of V1 interneurons to enter depolarization block with increasing stimulation (Figure 8A-8C, Figure S6A & see Methods). Using known connectivity and CSN input statistic approximations from *in vitro*^42,44^ and see Methods)) and our *in vivo* recordings, we first modelled the effect of spontaneous CSN input during baseline (CSN_Spont._, see Figure 3A) and found that under *in vivo*-like conditions the subthreshold membrane potential (V_m_) remained close to threshold resulting in a relatively high baseline firing rate (Figure 8D-8E). In this regard, spontaneous CSN input alone places V1 interneurons in a regime where they are sensitive to small fluctuations in V_m_, while large, sustained fluctuations result in output saturation and eventually depolarization block (Figure 8E and Figure S6B). Also, having a non-zero baseline firing rate facilitates bidirectional output modulation depending on changes in synaptic input patterns^46^. To investigate how changing CSN input might affect the input-output relationship during movement we considered three different input types. Time-locked, movement-related increased activity was considered as the driving input (CSN_Inc._), while spontaneous, background CSN activity (CSN_Spont._) (see Figure 3A) was classified as modulatory and could be adjusted to mimic the movement-related decrease in spontaneous activity observed in a subpopulation of CSNs *in vivo* (CSN_Dec._) (Figure 8F and Figure S6C-S6D).

**Figure 8.**
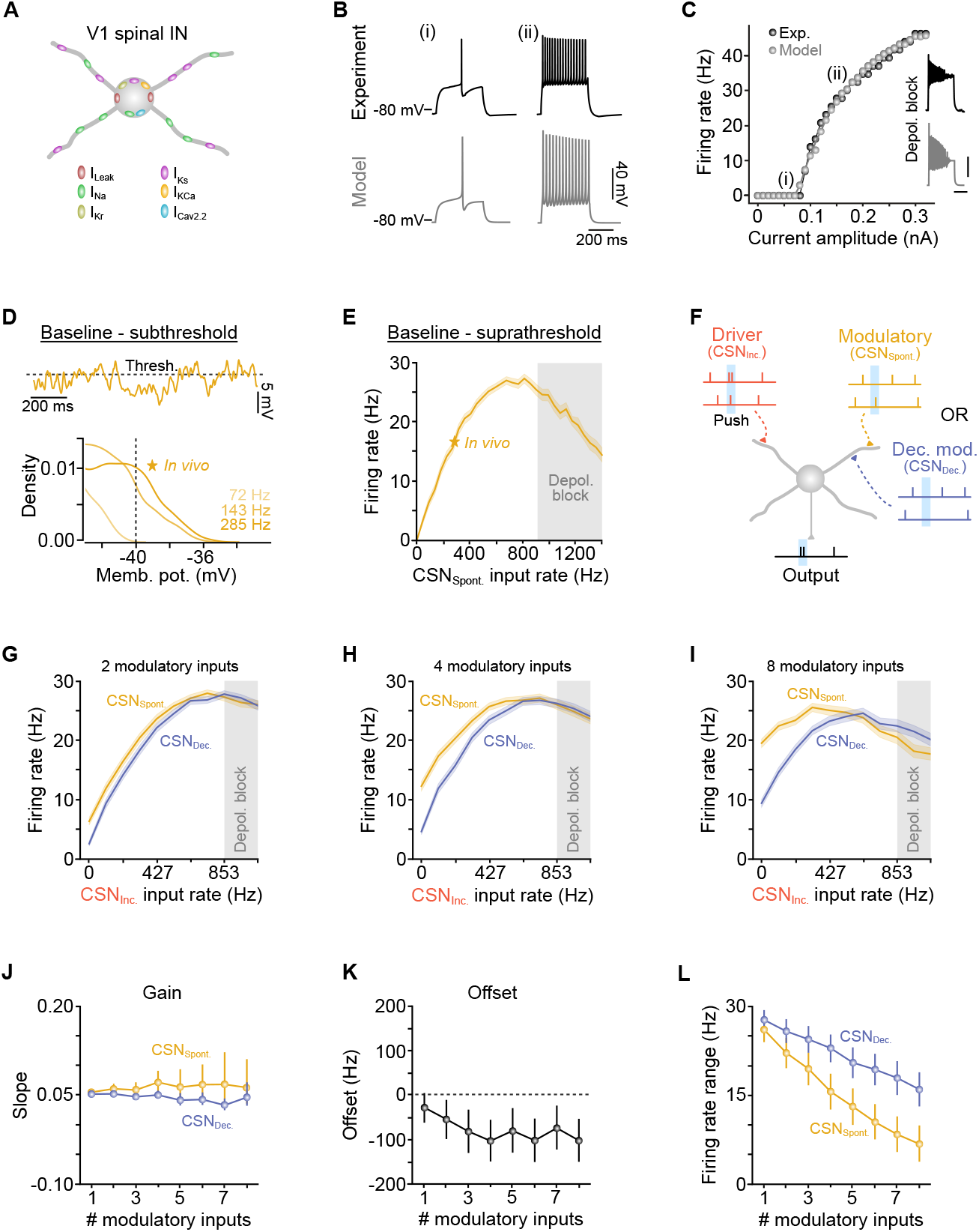
Reduced CSN input drives subtractive input-output transformations in V1 spinal interneurons. (A)Schematic of a V1 spinal interneuron model with leak (I_Leak_), fast sodium (I_Na_), delayed rectifier K+ (I_Kr_), slowly inactivating K+ (I _Ks_), Ca2+-dependent K+ (IK_Ca_) and N-type Ca2+(I_Cav2.2_) conductances. (B)Example voltage traces from an *in vitro* slice recording of an identified V1 spinal interneuron (black traces) and model equivalent (gray traces) during (i) 60 pA and (ii) 170 pA current injections. (C)Frequency-current (FI) relationship for V1 interneurons recoded *in vitro* (Exp., n = 7 cells) and model equivalent (Model). (i) and (ii) relate to the voltage traces shown in (B). Insets, show depolarisation block during current injection *in vitro* (*top*, 0.4 nA) and in the model (*bottom*, 0.8 nA). Scale bars, 40 mV, 200 ms. (D)*Top*, V1 model: baseline membrane potential with respect to threshold (dashed line) when modeling *in vivo*-like inputs. Model spiking mechanism has been removed to illustrate subthreshold dynamics. *Bottom*, all-point histogram of model membrane potential near to threshold for 72 Hz, 143 Hz and 285 Hz CSN input rates. (E)Model frequency-current (FI) relationship during baseline. Note moderate baseline firing rate in the presence of *in vivo*-like input statistics and tendency to enter depolarization block above 800 Hz input. (F)Schematic depicting coincident excitatory ‘driver’ input during lever pushes (CSN_Inc._, red) and spontaneous ‘modulatory’ input from layer 5B CSNs that do not change their firing rate during movement (CSN_Dec._, yellow). The effects of changing the modulatory input were modelled as a decrease in spontaneous input during movement (CSN_Spont._, blue) to mimic effects observed *in vivo*. (G-I) Model input-output relationships in the presence of 2 (G), 4 (H) and 8 (I) modulatory inputs (i.e. CSN _Spont._ Or CSN_Dec._). CSN_Dec._ input statistics derived from *in vivo* recordings. Gray shaded region, depolarization block. (J)Median slope of the input-output relationship as a function of increasing number of modulatory inputs (i.e. CSN_Spont._ or CSN_Dec._). (K)Median offset between CSN_Spont._ and CSN _Dec._ as a function of increasing number of modulatory inputs (i.e. CSN _Spont._ or CSN_Dec._). (L)Median firing rate range as a function of increasing number of modulatory inputs (i.e. CSN_Spont._ or CSN_Dec._).

This allowed us to directly test the effect of reducing the level of modulatory input on the input-output relationship of V1 interneurons and uncover any underlying arithmetic operations (Figure S6E-S6H)^47^. In the absence of decreased activity (CSN _Inc._ + CSN _Spont._), the output firing rate increased as a function of CSN input rate but became more susceptible to output saturation as the number of modulatory inputs was increased (Figure 8G-8I). This effect was due to the already depolarized baseline V_m_ being maintained close to or at threshold for longer periods, eventually leading to a breakdown of the spiking mechanism and depolarization block. In contrast, inclusion of a population of CSNs that decreased their activity (i.e. reduced modulatory input) (CSN_Inc._ + CSN_Dec._) resulted in a linear subtractive transformation of the V1 interneuron input-output relationship, without affecting the gain. This rightward shift (offset) was accompanied by a reduction in the probability of output saturation, and enhanced firing rate range (Figure 8G-8K) due to the reduction in background spontaneous excitatory input (i.e. modulatory input) driving the V_m_ away from threshold, which increased rheobase and decreased the postsynaptic effect of driving inputs. Overall, reduced CSN input enhanced the range across which driving inputs can influence V1 spike output before saturating, ensuring flexible input-output transformations across a range of modulatory input levels. Importantly, this linear additive operation is independent of the dorso-ventral location of CSNs in motor cortex but instead is shaped by the relative balance between driver and modulatory inputs. Given that multiple spinal interneuron subtypes receive CSN input from CFA^42^, this could in principle provide a canonical biophysical mechanism to ensure flexible input-output transformations across different spinal targets.

## Discussion

Reduced cortical output during behavior is common yet understudied, observed during sensory processing, decision-making, and motor control^1,17,24,48,49^. Despite its prevalence, its functional significance remains unclear. This is particularly striking in the motor system where reduced cortical output during movement is known to be conserved across species^1,15–18^, yet the focus has tended to be on projection neuron activation rather than suppression. Here, we addressed this by developing a Go/NoGo forelimb lever task for mice to show that reduced CSN output is selectively associated with skilled movement execution, is spatiotemporally organized, and drives subtractive modulation and flexible scaling of V1 spinal interneuron input-output transformations.

Previous studies in rodents have largely explored an activation-based model of CSN function shaped by the topographical organization of activity at different timepoints during movement^3^, projection target specificity^15^ and emerging patterns of activation across learning^1,4^. Data from our task align with this model, in that CSN activation strongly correlated with the magnitude of limb movements, rising before movement onset and returning to baseline upon movement completion^1,3,4,15^. Although widespread dendritic tuft activation dominated, indicating strong somatodendritic coupling^1,33,34^, we also observed marked functional compartmentalization within and between CSN tuft hemi-trees, consistent with parallel branch-specific computations being an important feature of ongoing motor control^31,36^. This activation model, however, overlooks the significant proportion of CSNs that reduce their activity during movement (8 - 35 % after motor learning)^1,15^, which in our task strongly correlated with the timing and magnitude of limb movements and dominated task-related activity (~75% of task modulated CSNs). The reason for the increased proportion of CSNs displaying reduced activity in our task is not clear, but is unlikely to be due to differences in recording location (somatic versus apical dendritic trunk) given the high degree of somato-dendritic coupling that we and others have observed in layer 5B projection neurons^1,33,34^. Instead, it may simply reflect differing task demands (i.e. constant torque multi-lever press versus single horizontal lever push)^1,15^, where the proportion of CSNs displaying reduced output scales with task-specific requirements. In this regard, the ratio and spatiotemporal organization of CSNs exhibiting increased versus decreased activity provides a flexible framework to support divergent CSN output patterns necessary for executing a wide range of movements with differing action specifications.

In rodents, skilled motor control correlates with reproducible spatiotemporal activity sequences in both superficial and deep layers of motor cortex^1,50,51^, with layer 2/3 exerting powerful top-down control of CSNs in layer 5B^30,52^. Importantly, only CSNs at the layer 5A/5B border (~200 µm region) receive strong descending input^52^, providing a functional architecture that is consistent with generating the dorsoventral activity gradient seen in our task. Whilst layer 2/3 provides the dominant input stream to upper layer 5B CSNs, inputs from secondary somatosensory cortex, motor thalamus, and layer 6 corticothalamic neurons may also contribute^28,53–55^. Given this input structure, what input source(s) might account for reduced CSN activity in deeper regions of M1 in our task? In principle, movement-related suppression of spontaneous input from orbitofrontal cortex, secondary motor cortex or motor thalamus could result in a lowering of the spontaneous event frequency and mean baseline ΔF/F_0_ ^28^. Alternatively, increased long-range excitatory input during movement could recruit local inhibitory interneurons in layer 5B providing strong feedforward inhibition that selectively dampens both spontaneous fluctuations and mean firing rate of deep layer CSNs^24,28,56–59^. Further studies, using a combination of cell-type- and pathway-specific recording and manipulation approaches, will be required to identify which input or combination of local and long-range inputs dictate the spatiotemporal organization of CSN activity during skilled movements.

Corticospinal neurons send collaterals to the basal ganglia, thalamus, midbrain, pons, and medulla en route to the spinal cord^19,20^, broadcasting information to supraspinal brain regions with localized specificity being endowed by differential inputoutput transformations at the level of the postsynaptic target^15^. Our finding that axons from both superficial and deeper CSN subpopulations converged across supraspinal areas except in the medulla where axon innervation is predominantly from deeper layer CSNs, is reminiscent of the projection-specific arrangement of PT neurons in the anteriolateral motor cortex (ALM)^41^. Given that CSN axon terminal densities in spinal cord far outweigh those in subcortical regions (~90% vs ~10%, respectively), the impact of collaterals via the cortico-brainstem-spinal cord pathway may only be modulatory in nature, with direct corticospinal input having the largest impact on spinal cord circuit dynamics^3^. The vast majority of CSN axons do not synapse on motor neurons but instead preferentially target inhibitory over excitatory pre-motor neurons, despite there being equal proportions of both neuronal classes in the mouse spinal cord^42,43^. Our anterograde and retrograde tracing suggested convergence of axons in the intermediate region of the spinal cord and on V1 interneurons, however, unequivocal confirmation would require retrograde rabies tracing from a single V1 interneuron starter cell, which is prohibitively challenging in the spinal cord. Instead, our dual mapping approach, coupled with previous CSN targeting studies suggest input convergence to be the most parsimonious explanation^2,3,44^.

Although we observed a clear dorsoventral gradient of CSN activity, this is likely to be specific to our task and we would expect different spatiotemporal activity patterns across a range of skilled actions such as lever press, pellet/water droplet reaching or joystick manipulation^1,4,60–62^. However, this depth-dependent functional architecture allowed us to investigate whether populations of CSNs with opposing dynamics converged at the level of the spinal cord. An important point to note is that downstream computations driven by bidirectional CSN input are agnostic to cortical location, instead they are shaped by the ratio of increased versus decreased CSN input. In this regard, reduced CSN input to V1 interneurons likely drives subtractive input-output changes irrespective of the movement type and spatiotemporal organization of upstream CSN activity. Although we did not record from CSNs deep in layer 5B (i.e. >750 µm), our model-based exploration of V1 input parameters and different ratios of increased versus decreased CSN inputs was designed to cover a wide range of possible CSN input combinations likely to occur *in vivo*.

How might subpopulations of CSNs with opposing dynamics influence downstream spinal cord circuit activity? By using a conductance-based V1 interneuron model, we found that under *in vivo*-like conditions spontaneous CSN input maintained the V_m_ close to threshold where stochastic fluctuations resulted in a relatively high baseline firing rate, consistent with recordings from cervical spinal interneurons in awake non-human primates^46^. Modelling movement related CSN activity suggested that reduced modulatory input (i.e., reduced spontaneous activity) generated a linear subtractive shift (offset) in the V1 input-output relationship, lowering the probability of output saturation and enhancing the output firing rate range. This combined effect is suggestive of a subtractive operation driven by both input and output modulation (Figure S8E and S8F), where the firing rate always remains non-zero given the continuous baseline input to CSNs. This subtractive shift could enable V1 interneurons to perform the same rate-coded input-output operation irrespective of the ratio of CSN_Inc._ and CSN ^47^, facilitating the linear transformation of CSN _Dec._ input to V1 output across a range of different activation patterns. Given the propensity of V1 interneurons to saturate and enter depolarization block with moderate input rates *in vitro* (see Figure 8C and Figure S6A), this mechanism could also provide a simple failsafe to avoid saturation and ensure a large dynamic output range. If left unchecked, V1 interneuron saturation and eventual depolarization block would disrupt input to downstream flexor and extensor motor neurons leading to disrupted muscle activation and limb discoordination^63,64^.

Given that there are anatomical and intrinsic electrophysiological similarities between differing subclasses of spinal interneurons receiving CSN input, it is inviting to speculate that the same biophysical mechanisms could generalize to other excitatory and inhibitory interneuron subtypes that reside in the intermediate region of the spinal cord^64–70^. What will be important to determine is how integration of different CSN input patterns is affected by subtle alterations in the complement of ion channels across interneuron subtypes, and to what extent these changes affect input-output transformations and arithmetic operations. Although we did not investigate the effect of local inhibition in our model, this additional type of modulatory input could in principle accentuate the subtractive modulation of the V1 input-output relationship seen in this study or drive divisive gain modulation depending on the nature of the movement-related inhibitory input^47,71–75^. An important next step will be to investigate how convergence of differing local and long range excitatory and inhibitory inputs contribute to shaping input-output transformations across different classes of spinal interneurons and their individual and combined contributions to skilled limb movement.

In summary, our findings highlight a critical role for reduced CSN output in shaping spinal cord computations during skilled movement execution, which raises important questions regarding downstream integration of cortical output. Could this be a generalized mechanism for shaping input-output transformations in supraspinal and brainstem targets receiving CSN input? Do these findings extend to other motor cortical projections neurons and their targets? How do direct and indirect signals from motor cortex combine at the level of the spinal cord and is reduced supraspinal input a consistent feature? Addressing these questions will be an important next step towards uncovering how motor cortical output shapes spinal cord computations necessary for performing goal-directed movements.

## Acknowledgments

We would like to thank members of the Duguid lab (Josh Dacre, Olivier Delree, Julia Schiemann, Federico Claudi and Julian Ammer) for help with designing the behavior, developing imaging approaches, image analysis and for experimental discussions throughout the project. We are grateful to Peter Rupprecht for assistance in adapting the CASCADE spike inference algorithm to accommodate our CSN data and Andre Rivard for conducting the *in vitro* slice recordings used to generate our V1 spinal interneuron model. AAV-GCaMP6s was a gift from Douglas Kim and the GENIE Project. AAV-pgk-Cre was a gift from Patrick Aebischer. pAAV-CAG-Flex-mRuby2-GSG-P2A-GCaMP6s-WPRE-pA was a gift from Tobias Bonho-<table> effer, Mark Huebener, and Tobias Rose. We would also like to thank Nathalie Rochefort and Valerio Francioni for advice on performing dendritic imaging and Tiago Branco and Kostas Betsios for their generous gift of the control software Mantis64. The research was supported by grants from the Biotechnology and Biological Sciences Research Council (BB/Y004639/1 to I.D. & M.S.R.), a Wellcome Senior Research Fellowship and Wellcome Discovery Award (110131/Z/15/Z & 227475/Z/23/Z to I.D.), National Natural Science Foundation of China (Grant No. 32192412 to X.L.) and STI2030-Major Projects (Grant No. 2021ZD0201002 to A.L.), National Institutes of Health (RO1NS047357 to F.J.A.; R01NS123116 to J.B.B.; U24EB028998, P50MH109429, R01MH140020 and R56MH132637 to S.D-B.), American Lebanese Syrian Associated Charities (J.B.B.), New York State Department of Health (DOH01-C38328GG to S.D-B.).

## Methods

### Experimental models and subject details

Male and female adult C57BL/6J&N, wild-type and Ai14(RCL-tdT)-D mice (The Jackson Laboratory strain #:007914, USA) (5-12 weeks old, 20-30 g, 1-4 animals per cage) were maintained on a reversed 12:12 hour light:dark cycle (lights off at 7 am) while *En1*::*Cre*;*Foxp2::Flpo* mice of both sexes (8 weeks old) were maintained on a standard 12:12 hour light:dark cycle. All animals were provided with *ad libitum* access to food / water and environmental cage enrichment (e.g., cardboard tubes, plastic domes, chewing sticks, and rope ladders). Experiments and procedures were approved by The University of Edinburgh local ethical review committee and performed under license from the UK Home Office in accordance with the Animal (Scientific Procedures) Act 1986 (Duguid), or the Institutional Animal Care and Use Committee of St. Jude Children’s Research Hospital and performed in accordance with the National Institutes of Health (NIH) guidelines.

## Contributions Matrix

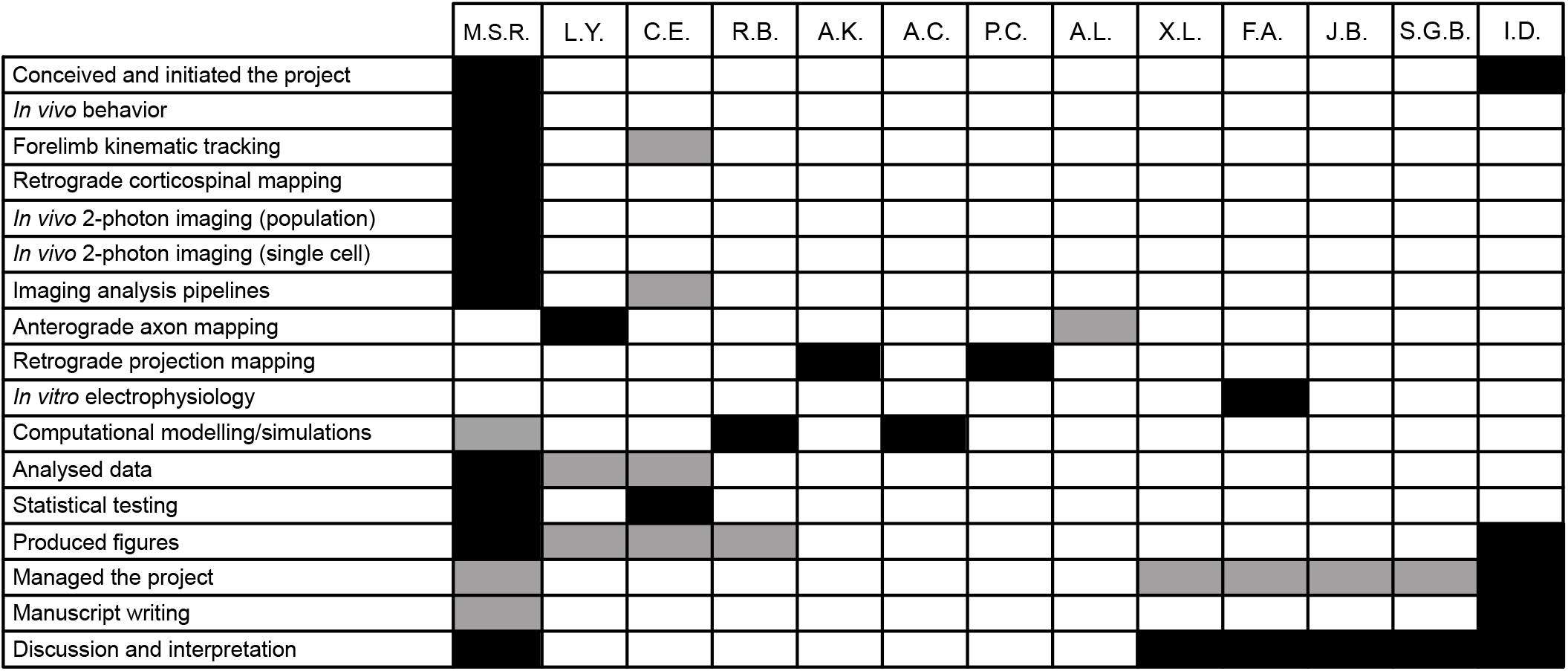

### General surgery and headplate attachment for imaging

Surgical procedures were performed under ~1.5% isoflurane anesthesia and each animal received fluid replacement therapy (0.5 ml sterile Ringer’s solution), buprenorphine (0.1 mg/kg), carprofen (20 mg/kg) and dexamethasone (2 mg/kg) for pain relief and to reduce inflammation. Mice were placed in a stereotactic frame (David Kopf Instruments, CA, USA) on top of an isothermal heating blanket with core body temperature maintained at 37oC.

In mice undergoing head-fixed behavioural training, the scalp was removed and the edges of the skin were attached to the exposed skull using Vetbond Tissue Adhesive (3M). A small RIVETS 3D printed headplate (0.5 g)^76^ was implanted on the surface of the skull using cyanoacrylate glue and dental cement (Lang Dental, IL, USA).

After surgery, mice were placed in a temperature-controlled recovery cage at 28oC to recover from the effects of anesthesia before being returned to their home cage. For post-operative care, Carprofen (20 mg/kg) was administered subcutaneously at 24 and 48 hours after each surgery.

### Go/NoGo lever push task design

The task design required mice to rest their right forepaw on a stationary lever and left forepaw on a moveable lever where the ‘reset’ mechanism was controlled by a servo motor driven by a Picaxe-08 Servo Driver board. The lever position (either ‘back’ or ‘front’) was determined by infra-red (IR) beams placed at both ends of the lever travel (4 mm). Each trial began with a 6 s inter-trial interval (ITI) followed by presentation of an auditory cue (240 ms long, Go: 6 kHz or NoGo: 13kHz, delivered in a pseudo-randomized order), and then a 4 s response window. Successful Go trials (i.e. lever push) were classified as ‘Hits’ and were rewarded with a 5 µl water droplet delivered 1 s after completion of the lever push. Water was delivered through a movable spout and controlled by a solenoid valve. The spout remained in place for 2 s to ensure complete reward consumption and the lever was reset and locked in the ‘back’ position during this period. Unsuccessful Go trials (i.e. no lever push), classified as ‘Miss’ trials, were neither rewarded nor penalized and the end of the response window was signaled by the lever being reset to the ‘back’ position and locked for 1 s. Successful NoGo trials (i.e. release and re-grasps) were classified as ‘Correct Rejections’ (CRs) and were rewarded at the end of the 4 s response window. Unsuccessful NoGo trials (i.e. lever push), classified as ‘False Alarms’, evoked a 80 ms burst of white noise followed by a 7.5 s time-out period during which the lever was reset and locked in the ‘back’ position (Fig S1A). Incomplete lever pushes during FAs were neither rewarded nor penalized and the end of the response window was signaled by the lever being reset to the ‘back’ position and locked for 1 s. Spontaneous lever pushes during the ITI were penalized with a 80 ms burst of white noise immediately after the lever left the ‘back’ position, followed by a 7.5 s time-out period during which the lever was reset and locked in the ‘back’ position. An Arduino Uno control board running custom written code monitored the status of the IR sensors, controlled the servos and solenoid, and generated the auditory cues. Signals from the Arduino were recorded as voltage changes at a sampling rate of 1 KHz via a data acquisition device (USB-201, Measurement ComputingTM) coupled to the Arduino board.

### Behavioral training

To enhance task engagement, mice were placed on a water control paradigm (1 ml / day) 7 days prior to training and weighed daily to ensure body weight remained above 85% of baseline. During this period, mice were handled extensively and habituated to the head restraint apparatus. Once habituated, mice were encouraged to interact with the moveable lever in a single 30 min session and rewarded for executing self-initiated lever pushes up to a maximum of 100 rewards. Mice were then trained on the Go/NoGo task once per day for 40 mins, starting with a 2-day ‘shaping’ phase. On the first ‘shaping’ day the duration of the auditory cues was set to 4 s, and then reduced to 2 s on the second ‘shaping’ day. Mice achieving a threshold of 50 rewards on two consecutive ‘shaping’ days progressed to the ‘training’ phase.

Performance was evaluated using the discriminability measure (*d’*) from the inverse normal cumulative distribution of the Hit and FA rates: *d’* = z(Hit) − z(FA). To account for the effects of satiation on motivation and performance^25^, d’ was calculated using a rolling window analysis (10 min duration, 2 min step size). Mice who maintained a *d’* > 1.5 for at least 50% of the session across three consecutive sessions were considered to have reached ‘expert’ level. Correction trials, in which an unsuccessful trial was followed by the same trial type until the correct action was performed, were present throughout training until ‘expert’ level had been attained. Correction trials were not used during recording sessions.

### Forelimb kinematic tracking

Mouse forelimb kinematics were recorded using a moderate frame rate camera (50 fps, Mako U U-029, AlliedVision) and controlled by Mantis64 (https://www.mantis64.com/). To measure gross forelimb movements, a region of interest (ROI) was manually drawn around the left forelimb and the frame-to-frame difference in pixel intensity was calculated using the formula:

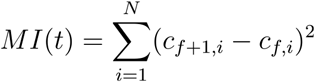

where c*f,i* is the grayscale level of pixel *i* in frame *f*. This generated a ‘motion index’ (MI). Aligning the motion index to cue presentation allowed identification of MI onsets (i.e. the first point where MI > threshold set as 2.5 × standard deviation of the motion index during 1 s baseline prior to cue). Movement correlations were calculated as the pairwise Pearson’s correlation coefficient of the MI during the 500 ms following cue presentation.

Vector tracking of the forelimb and lever movement was performed using DeepLabCut^77^. Tracking data (X and Y coordinates) were aligned to cue and baselined to mean position during the 100 ms prior to cue presentation. Movement vectors were calculated from the trial-averaged paw position at cue and at median onset of the lever push.

Spontaneous release and regrasps were identified by finding all instances of forelimb movements during the ITIs (z-scored MI > 1.8) and using vector tracking data to sub select those in which the direction of the corresponding movement vector was within 45° of the average cued release and regrasp vector for each mouse.

### Retrograde tracing of CSNs

To selectively label corticospinal neurons (CSNs) in CFA, we injected a retrograde adeno-associated virus (AAV) AAV-pgk-Cre (1.7×10^13^ vg/ml, Addgene viral prep #24593-AAVrg) into the spinal cord of Ai14(RCL-tdT)-D mice. To gain access to the spinal cord, an incision was made in the skin from the top of the neck to the bottom of the shoulder blades. Subcutaneous fatty tissue was removed, exposing the trapezius muscles. The muscle fibers were carefully separated along the midline using blunt forceps and held apart with retractors to expose the spine. A partial laminectomy was performed at C6 exposing only the left side of the spinal cord. 50-80 nL of the virus were injected (two sites: 0.3 mm lateral to the midline, 0.6 mm and 0.3 mm ventral to the surface). Virus was delivered via a pulled glass pipette (5μl, Drummond Scientific, PA, USA) using an automated injection system (Model Picospritzer iii, Parker, NH, USA; 10–20 nL/min). After injection, the trapezius muscles were gently pushed back in place with a pair of cotton buds, and the wound was closed using 5-0 Vicryl sutures (Ethicon, UK).

After 14 days, mice were terminally anaesthetized using an intraperitoneal injection of sodium pentobarbital (Euthatal, Merial, 0.10–0.15 ml) and transcardially perfused with 30 ml of ice-cold 0.1 M phosphate-buffered saline (PBS) followed by 30 ml of 4% paraformaldehyde (PFA, Sigma-Aldrich, MO, USA) in 0.1 M PBS solution and adjusted to pH 7.4. Brains were post-fixed at 4°C for 1–3 days in 4% PFA solution, then transferred to PBS solution. Individual brains were cut into parasagittal sections (60 μm) using a vibrating microtome (Leica VT1200S, Leica Microsystems (UK) Ltd.), counterstained using a green fluorescent Nissl stain (NeuroTrace 500/525, ThermoFisher) and mounted with Vectashield Antifade Mounting Medium (Vector Laboratories, CA, USA). Images were acquired with a Leica DM R epifluorescence microscope and image analysis was performed using Fiji (National Institutes of Health; https://imagej.net/software/fiji) and MATLAB (MathWorks, MA, USA). To generate a spatial map of C6-CSNs, images were aligned to the Paxinos mouse brain atlas^78^ to identify the medio-lateral (ML) location of each brain slice relative to bregma. Then, slices were divided into 100 μm bins in the antero-posterior (AP) axis. Td-Tomato expressing CSNs were counted in each bin of every second slice using the Fiji Particle Analysis toolbox. To facilitate automatic cell-counting, images were pre-processed using background subtraction (rolling ball, 15 pixels wide at 1.28 μm/pixel) and binarized using a thresholding method. Overlapping cell bodies were separated using a water shedding algorithm and filtered based on size (>100 pixel area) and circularity (0.5 to 1). To construct a depth profile of CSNs, the binarized cortical sections were divided into bins 25 μm deep. The amount of fluorescence in each bin was calculated as the number of bright pixels (pixels with a value of 1), normalized within slice, and averaged across slices.

### 2-photon calcium imaging

To selectively express the genetically encoded calcium indicator GCaMP6^79^ in CSNs that project to the C6 segment of the spinal cord, we used a similar intersectional viral strategy as described previously, with some modifications. To gain access to the C6 spinal cord, an incision was made in the skin from the top of the neck to the bottom of the shoulder blades. Subcutaneous fatty tissue was removed, exposing the trapezius muscles. The muscle fibers were carefully separated along the midline using blunt forceps and held apart with retractors to expose the spine. A partial laminectomy was performed above C6 exposing only the left side of the spinal cord, ipsilateral to the moving limb. 50-80 nL of AAVrg-pgk-Cre (1.7×10^13^ vg/ml, Addgene viral prep #24593-AAVrg) were injected into the spinal cord (two sites: 0.3 mm lateral to the midline, 0.6 mm and 0.3 mm ventral to the surface). Virus was delivered via a pulled glass pipette (5μl, Drummond Scientific, PA, USA) using an automated injection system (Model Picospritzer iii, Parker, NH, USA; 10–20 nL/min). After injection, the trapezius muscles were gently pushed back in place with a pair of cotton buds, and the wound was closed using 5-0 Vicryl sutures (Ethicon, UK). Cortical injections and headplate attachment were performed 4-7 days after spinal injections. The scalp was removed, and the edges of the skin were attached to the exposed skull using Vetbond Tissue Adhesive (3M). A small burrhole was made using a hand-held dentist drill with 0.5 mm burr and 250 nL of AAV-Syn-FLEX-GCaMP6f (diluted to 1.7×10^12^ vg/ml, Addgene 100833-AAV1) was injected into CFA (1.6 mm lateral, 0.2 mm rostral to bregma and 0.7 mm from the cortical surface). Following injection, the craniotomy was sealed with a small volume of platinum-cure silicone (Body Double Fast Set, Smooth-On). Finally, a small lightweight RIVETS headplate (0.5 g) was implanted on the surface of the skull using cyanoacrylate glue and dental cement (Lang Dental, IL, USA). After behavioural training, a 3 mm ø circular cranial window (glass coverslip #0; Menzel-Gläser, Germany) was implanted above the virus injection site and fixed to the skull with cyanoacrylate glue.

For dendritic calcium imaging, we used the same intersectional viral strategy with the following modifications: (1) to achieve sparse labelling, AAVrg-pgk-Cre (Addgene viral prep #24593-AAVrg) diluted 1:150 was injected into the C6 segment of the spinal cord (two sites: 0.3 mm lateral to the midline, 0.6 mm and 0.3 mm ventral to the surface), and (2) we introduced a calcium-independent structural marker by injecting 250 nL of a bicistronic viral vector expressing GCaMP6s and the fluorescent protein mRuby2^32^ into CFA (AAV-CAG-Flex-mRuby2-GSG-P2A-GCaMP6s, 1.7×10^12^ vg/ml, Addgene viral prep #68717-AAV1) into CFA (1.6 mm lateral, 0.2 mm rostral to bregma and 0.7 mm from the cortical surface). Cranial window implantation surgery involved placing a triple glass window consisting of two 3 mm ø circular coverslips glued under a 4 mm ø coverslip, such that the two small glass discs were inserted into the craniotomy while the larger coverslip provided a fixation point to the skull. This minimized axial brain movement sufficient to image small diameter dendrites.

2-photon calcium imaging of the somata of CSNs was performed using an 8 kHz resonant scanning microscope (HyperScope, Scientifica, UK; 690 × 690 μm FOV; 512 × 512 pixels) with a Ti:Sapphire pulsed laser (Chameleon Vision-S, Coherent, CA, USA; < 75 fs pulse width, 80 MHz repetition rate) tuned to 920 nm. Images were acquired at ~30 Hz with a 16x objective lens (0.8 NA; Nikon) using SciScan image software (Scientifica) and synchronized with high-speed video and behavioral data acquisition using Mantis64. Dendritic imaging was performed using a custom-built resonant scanning 2-photon microscope (320 × 320 μm FOV; 600 × 600 pixels) with a Ti:Sapphire pulsed laser (Chameleon Vision-S, Coherent, CA, USA; < 75 fs pulse width, 80 MHz repetition rate) tuned to 920 nm wavelength. Images were acquired at 50 Hz with a 40x objective lens (0.8 NA; Nikon) and custom-programmed Lab-VIEW-based software (LotoScan). Near-simultaneous multiplane imaging was performed by mounting the objective on a Piezo objective positioner (P725.4CA, Physik Instruments, Germany). Two planes 90 - 210 μm were selected, resulting in an objective travel time of 40 ms between planes, with a sampling rate of 8.33 Hz. Most CSN dendrites were imaged across 2 days, resulting in 4 separate imaging planes.

Motion artifacts in the raw fluorescence videos were corrected using discrete Fourier 2-dimensional-based image alignment (SIMA 1.3.2^80^). ROIs were drawn manually in Fiji and pixel intensity within each ROI averaged to produce a raw fluorescence time series (F). To remove fluorescence originating from neuropil and neighboring neurons, fluorescence signals were decontaminated and extracted using nonnegative matrix factorization, as implemented in FISSA^81^. Normalized fluorescence was calculated as ΔF/F_0_, where F _0_was calculated as the 5^th^ percentile of the 1 Hz low-pass filtered raw fluorescence signal and ΔF = F-F_0_. All further analyses were performed in custom written scripts in MATLAB or Python 3.

### Functional cell-type classification

To identify CSNs with somatic movement-related activity, we defined a baseline as the 1 s period prior to cue presentation and a ‘response’ epoch as the 1s after cue presentation. The response epoch was divided into 250 ms rolling time bins (50 ms step size), and the average ΔF/F_0_ of each bin was compared to the average ΔF/F_0_ of the baseline epoch. A boot-strapped distribution (10,000 samples) of the difference ΔF/F _0(bin)_ - ΔF/F _0(baseline)_ was obtained across trials and used to test whether the 99% confidence intervals were different from 0. Neurons were classified as movement-related if: (1) they contained 3 or more consecutive bins significantly different from baseline, and (2) the onset of the change (see ‘*Identification of activity change onset and offset times*’ below) was detected between cue and median push completion.

To identify CSNs with dendritic movement-related activity, we first obtained the average ΔF/F_0_ across all branches. Due to the lower acquisition rate, we could not use a rolling time bin approach so instead we compared the mean ΔF/F_0_ in a baseline epoch 1 s prior to cue to the mean ΔF/F^0^ in a single response epoch 500 ms following cue. A bootstrapped distribution (10,000 samples) of the difference ΔF/F_0(response)_ - ΔF/F _0(baseline)_ was obtained across trials and used to test whether the 99% confidence intervals were different from 0. This approach minimized the likelihood of including events with onsets towards the end of the response window.

Activity rasters of movement-related activity are presented as trial averages, baselined to the average ΔF/F_0_ in the 1 s precue period, and normalized to the maximum deviation from baseline (whether positive or negative). Push trials (Hits and False Alarms) are shown aligned to the onset of the lever movement. Release and regrasp trials (Correct Rejections) are shown aligned to the median lever movement onset during Hit trials.

### Spike inference from calcium imaging data

Calcium signals were deconvolved using the CASCADE algorithm^26^, consisting of a deep network trained on noise- and rate-matched ground truth datasets. The algorithm utilizes unsmoothed ΔF/F^0^ traces to estimate the expected number of spikes in each frame. This ‘spike probability’ was then multiplied by the recording frame rate to obtain an estimate of the instantaneous firing rate for each cell.

To ensure high performance, we first used the CASCADE toolbox to estimate the standardized noise level in the fluorescence signal for each of our ROIs. Noise was calculated as

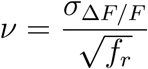

where *σ*ΔF/F is the median frame-to-frame difference in ΔF/F_0_ values and *f*_*r*_ is the recording frame rate.

For deep, somatic recordings in which noise levels were outside the range of pre-trained models, a new model (Global_ EXC_30Hz_smoothing100ms_causalkernel_high_noise) was trained with a matched range of noise levels.

### Identification of onset and offset times

The median onset times for movement-related CSN activity were calculated using the inferred spike rate and slope sum function onset detection algorithm (SSF^23,24,27^). The SSF uses a rolling window to enhance the upslope of spike rate traces (SR) through the equation:

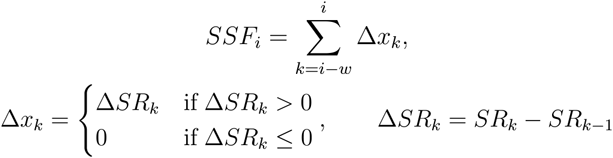

To identify onsets, we calculated the SSF using a window *w* = 1 frame. We identified the peak SSF after each cue presentation, and then performed a backwards search until the first point exceeding a threshold (10% of peak) was found. For neurons with decreased activity, the slope sum function was modified to identify negative (rather than positive) slopes and then inverted (multiplied by −1) so that the peak change and onset could be detected in the same way as increased activity. Onsets are reported as the median of 10,000 bootstrapped samples to reduce the influence of noisy individual trials. To detect any systematic biases in our onset detection and to calculate a correction factor, we verified the algorithm with ‘ground truth’ datasets available at https://github.com/HelmchenLabSoftware/Cascade. The ground truth ΔF/F_0_ signals were deconvolved with the same model as our data, then detected onsets were compared to the times at which the corresponding spike trains started or ended. This validation step was performed for both subpopulations of CSNs, resulting in a correction factor of −32 ms, and +13 ms for increased and decreased activity, respectively.

### Correlation between activity and movement

To investigate the relationship between neuronal activity and forelimb movement, the correlation coefficient (Pearson’s r) was calculated between the trial-averaged forelimb motion index (MI) and spike rate of increased and decreased CSN activity in a 1 s peri-movement window. MI and spike rate time series were smoothed with a 0.5 s LOESS filter and then the bootstrapped median was calculated for each cell by creating 10,000 bootstrap samples of 10 randomly selected trials (sampled with repetition). To ensure statistical robustness, animals with less than 5 CSNs (increased or decreased) were excluded from analysis and results are presented as the median per animal.

### Mapping dendritic activity

Dendritic morphologies were reconstructed through ultra-sparse labelling of CSNs. At the end of each imaging session, z-stacks of mRuby2 fluorescence (laser tuned to 1020 nm) were acquired at 1 μm steps in the axial plane. The Fiji plugin Simple Neurite Tracer (SNT) was used to semi-automatically trace apical and tuft dendrites to generate a dendrogram and identify the somatic depth of each neuron. Imaged ROIs were aligned to the dendritic reconstruction and manually mapped onto the reconstructed dendrogram. The SNT function *measure* was used to obtain the length of each dendritic ‘segment’, where a ‘segment’ was defined as a portion of dendrite between branching points. The nexus length was calculated as the summed distance of the two dendritic segments between the first and second branching points^31^. Hemi-trees were defined as the two groups of branches separated by the first branching point.

Dendritic events were identified from z-scored ΔF/F_0_ time series using the Matlab function *findpeaks* with a minimum peak prominence of 1. The onset of each event was defined as the timepoint preceding the peak at which ΔF/F^0^ > 10% of peak. In each trial, a dendrite was defined as ‘active’ if an event occurred with an onset between cue and lever push completion (or median lever push completion in the case of release and re-grasps). For each sampled dendrite, the probability of activation during the movement period was compared to the probability of activation during the baseline period to generate a spatial map of increased or decreased activity throughout the dendritic tree. Movement related activation patterns during each trial were classified as: ‘whole tree’, trials where dendrites in both hemi-trees increased or decreased their activity; ‘hemi-tree’, where activity changes happened exclusively in one of the two hemi-trees (Main hemi-tree with the other being classified as Secondary hemi-tree); ‘single branch’, one isolated dendritic segment increased or decreased its activity; and ‘None’, no dendritic segments increased or decreased their activity.

### CSN axon reconstructions / fMOST imaging

Mice were anaesthetized with 1.2% tribromoethanol (Nanjing Aibei Biotechnology Co., Ltd., Nanjing, China) and C6-projecting CSNs were sparsely labelled by injecting 200 nL of AAVrg-hSyn-CRE (5×10^12^ vg/ml, Brain Case) into the C6 segment of the spinal cord and 200 nL of AAV9-EF1a-DIO-EGFP (3.5×10^12^ vg/ml) into the CFA of mice, as previously described. After 4 weeks, mice were anesthetized with a sodium pentobarbital solution (100 g/mL) and intracardially perfused with 0.01M PBS (Sigma-Aldrich) followed by 4% PFA (Sigma-Aldrich). The brain and spinal cord were removed and post-fixed in a 4% PFA solution at 4°C for 12 hours. After fixation, brains were dehydrated in a graded series of ethanol solutions (50%, 75%, 95%, and 100%; 4°C, 1 hour each). The dehydrated brains were then immersed in a graded series of Lowicryl HM20 resin (Electron Microscopy Sciences, China) (50%, 75%, and 100%; 4°C, 2 hours each) at 4°C for 2 hours at each concentration. Finally, the samples were impregnated with 100% HM20 resin for 2 days at 4°C and then embedded in 100% HM20 with 0.7% divinylbenzene at 20°C for 48 hours.

To save sample preparation space and imaging time, the spinal cord was wrapped around the brain. Specifically, it was bent from the medulla to the ventral side of the brain towards the olfactory bulb, and then returned along the dorsal side.

Resin-embedded brain and spinal cord preparations were imaged using an HD-fMOST system^38^. Fluorescence signals from illuminated samples were collected using a microscope objective (1.0 NA, XLUMPLFLN 20XW, Olympus) and projected through a dichroic mirror (ZT488/561rpc, Chroma) and an emission filter (ZET488/561m, Chroma) onto the light-sensitive surface of a scientific complementary metal oxide semiconductor (sCMOS) detector (ORCA-Flash 4.0, Hamamatsu). The surface of the sample was imaged in the coronal plane using an x-axis strip scanning pattern combined with y-axis splicing and a 1 μm z step resulting in a 3D image. The sample was then sliced with a diamond knife at a thickness of 2 μm. In total, the whole sample was imaged at a voxel resolution of 0.32 × 0.32 × 1 μm over ~72 hours.

The morphology of individual CSNs was semi-automatically reconstructed using GTree (https://github.com/GTreeSoftware/GTree). Using the Allen Mouse Brain Common Coordinate Framework version 3 (Allen CCFv3) brain atlas as a reference, the dataset was rotated and resampled to a voxel resolution of 10 × 10 × 10 μm^3^. Fluorescence images were registered to CCFv3 templates using the Symmetric Normalization (SyN) algorithm. Finally, the full dataset was imported into Amira Software (v6.1.1, FEI) and aligned with the 3D outline of corresponding brain regions.

Reconstructed CSNs were classified as ‘superficial’ if their soma was located at a cortical depth ≦ 650 μm, or ‘deep’ if located > 650 μm. The structure of each reconstructed neuron consisted of a continuous series of connected nodes which represented the spatial coordinates that were aligned to the CCFv3. The end of each axon branch or segment was defined as the axon terminal and counted for each segmented brain region.

To analyze the distribution of axons in the spinal cord, a series of 100 μm maximum intensity projections was generated from cervical spinal sections. Spinal cord segments C1-C8 were identified using visual landmarks according to Watson & Gulgun, 2009^82^. To compensate for the inaccuracies of identifying specific boundaries between spinal sections, data from sections C5-C7 were included. Each of the resulting fluorescence images was aligned to the SpinalJ C6 template (https://zenodo.org/records/5177185), via an affine transformation to correct distortions induced by the curvature of the tissue during imaging. Images containing isolated reconstructed axons were processed and transformed according to its corresponding fluorescence image.

Axon overlap in the spinal cord was mapped using maximum intensity projections from ‘superficial’ and ‘deeper’ CSNs. This resulted in two binary images in which the pixel color (white or black) depicted the presence or absence of a reconstructed axonal segment in that location. A third binary image was generated by identifying the pixels that overlapped and converted to a heatmap by smoothing with a Gaussian filter (σ = 30 pixels) and false-coloring with jet colormap.

### Labelling V1-projecting CSNs

V1-projecting CSNs were identified from serial two-photon tomography (STPT) images of rabies virus transsynaptic tracing experiments previously described in (Chapman et al., 2025)^44^. Briefly, Cre dependent helper viruses AAV-FLEX-TVA-mCherry (6.0 × 10^12^ vg/mL, UNC Vector Core) and AAV-FLEX-H2B-HA-N2cG (1.0 × 10^13^ vg/mL, Vigene Biosciences) were injected into C6-C8 segments of the spinal cord of P14 *En1::Cre* mice, as previously described. 5 weeks after the first injection, RV-CVS-N2cΔG-H2B-eGFP (3.5 × 10^6^ infectious units/mL) was injected into C6-C8 spinal segments. To label V1^Foxp2^-projecting CSNs, the Cre and Flp dependent helper viruses AAV-Con/Fon-TVA-mCherry (1.1 × 10^12^ vg/mL, Stanford Gene Vector and Virus Core) and AAV-Con/Fon-N2cG (3.1 × 10^12^ vg/mL, Stanford Gene Vector and Virus Core) were injected into C6-C8 segments of the spinal cord of P14 *En1::Cre; Foxp2::Flpo* mice, followed 5 weeks later by injection of RV-CVS-N2cΔG-H2B-eGFP (3.5 × 10^6^ infectious units/ mL), as above.

For both V1- and V1^Foxp2^-projecting CSN labelling, 7 days after the second injection, mice were terminally anaesthetized with Avertin (Tribromoethanol, 240 mg/kg body weight) and transcardially perfused with phosphate-buffered saline (PBS) followed by ice-cold 4% paraformaldehyde (PFA, Sigma-Aldrich, MO, USA) in 0.1 M PBS solution. Brains were postfixed at 4°C for 24 hours in 4% PFA solution and imaged via STPT on a high-speed multiphoton microscope with integrated vibratome sectioning (TissueCyte 1000, TissueVision, 16x/0.8 NA objective). The first images were acquired at 25, 50, and 75 μm below the surface at a resolution of 0.875 μm /pixel in X-Y plane at a Z-plane sampling interval of 25 μm, before a 75 µm coronal section was cut using a vibrating microtome. This image-cut sequence was then repeated until the whole brain had been imaged.

To visualize the distribution of V1-projecting CSNs in CFA a maximum intensity projection was created from all sections spanning +1 mm to −1mm antero-posterior to bregma. The resulting images were smoothed using a Gaussian filter (σ = 55 pixels), registered to each other (using a simple translation) and averaged. This across-animal average image was smoothed with a Gaussian filter (σ = 100 pixels) and cropped to a small region centered on CFA. The cropped image was thresholded at the 99.5^th^ percentile, false-colored with a jet colormap, and transformed to a contour map (Matplotlib function ‘contour’).

To construct a depth (dorso-ventral) profile of V1-projecting CSNs, the maximum intensity projection of each animal was cropped around CFA, and then thresholded at the 90^th^ percentile to facilitate using pixel intensity to quantify the presence of labelled CSNs. For each image, pixel intensities were averaged across the medio-lateral axis, resulting in a dorso-ventral profile, and then averaged across animals.

A depth profile of V1^Foxp2^-projecting CSNs was created by manually annotating the X and Y pixel coordinates of the center of each labelled soma in sections corresponding to antero-posterior locations ranging from +1 mm to −1 mm from bregma. Using the OpenCV library, a contour was created around each brain slice (function *findContours*), and the cortical depth of each neuron was calculated as the shortest distance between the soma and the contour (function *pointPolygonTest*). To construct a depth profile of V1^Foxp2^-projecting CSNs, cortical depths were divided in 50 μm bins, and cell density calculated as the number of cells counted per bin, then normalized within mouse and averaged across mice.

### Statistics

Data were analyzed using custom scripts written in Python (v3.12) and MATLAB.

#### Bootstrapped median differences for paired data

To determine whether the onsets and trial-to-trial correlations of the motion index (MI) differed across trial types (Hits, FAs and CRs), 10,000 bootstrap replicates were generated for each pair of trial types. In each replicate, existing animals were randomly selected with replacement to create a bootstrap sample of size n = 50. Then, the difference between the across-animals median onset or trial-to-trial correlations in the two different trial types (Δ _T1T2_) was calculated. After repeating this process 10,000 times, the resulting distribution of Δ_T1T2_ values was visually inspected to determine whether the bootstrap had converged (i.e. Δ_T1T2_ values were normally distributed^83^. Bootstraps which did not converge were repeated with 50,000 replicates and a sample size of 150 animals per replicate. Confidence intervals (95% CI) were drawn from the boot-strapped distribution using the percentile method^84^. p-values were derived from the 95% CI through the formula:

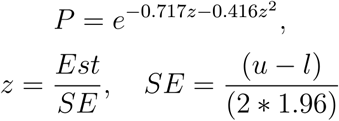

where *u* and *l* are the upper and lower bounds of the CI and *Est* is the estimated difference between groups^85^.

#### Hierarchical bootstrap for median differences

To compare the onset times and baseline activity levels between CSNs that displayed increased versus decreased activity, we utilized a hierarchical bootstrap method described by Ren & Saravanan^86,87^. To account for the variability in the proportion of increased and decreased CSNs across animals (FOVs), the hierarchical bootstrap was constructed to resample both CSN populations within animals as well as between animals. For each comparison, 10,000 ‘level 1 bootstrap replicates’ were constructed by resampling animals 50 times with replacement in each replicate. Within each resampled animal, CSNs displaying increased versus decreased activity were also resampled 50 times with replacement to obtain the difference between the median of the onset times / baseline activity levels of increased and decreased CSNs, constituting a ‘level 2 bootstrap replicate’. This process was repeated 10,000 times to create a ‘level 2 bootstrap sample’, and the mean of this distribution was taken as the ‘level 1 bootstrap sample’ for that replicate. Finally, confidence intervals (95% CI) and were drawn from the distribution of ‘level 1 bootstrap replicates’ and p-values derived as described in the previous section.

### In-vitro recordings of V1 spinal interneurons

Transverse spinal cord slices (300 µm thick) were prepared from postnatal day 10–14 mice to investigate the passive and active properties of V1 interneurons. Following rapid dissection in ice-cold, oxygenated (95% O_2_ / 5% CO_2_) sucrose-based artificial cerebrospinal fluid (aCSF; composition in mM: 218 sucrose, 26 NaHCO_3_, 3 KCl, 1.25 NaH_2_PO_4_, 10 D-glucose, 2 MgCl_2_), the tissue was embedded in low-melting agarose. Slices were sectioned using a Leica VT1000P vibratome before being sequentially transferred to an incubation chamber continuously perfused with oxygenated standard aCSF (in mM: 133 NaCl, 26 NaHCO_3_, 3 KCl, 1.25 NaH_2_PO_4_, 10 D-glucose, 2 MgCl_2_, 2 CaCl_2_). Slices were initially incubated at 32°C for 1 hr before being maintained at room temperature (22–25°C) until recordings.

For recordings, individual slices were transferred to a submerged chamber perfused with oxygenated aCSF at room temperature. The passive and active properties of V1^FoxP2^ interneurons were recorded in slices from *En1::Cre*; *FoxP2:*

*:Flpo*; RCE.dual.GFP mice which express GFP in V1^FoxP2^ interneurons as described in Bikoff *et al*., 2016^45^ using epifluorescence microscopy (Olympus BX50WI, 40X water-immersion objective). Patch pipettes were filled with an internal solution (in mM: 122.5 potassium D-gluconate, 17.5 KCl, 9 NaCl, 1 MgCl_2_, 10 HEPES, 3 Mg-ATP, 0.3 Na-GTP, 0.1% Cascade Blue; pH 7.2–7.3), with 20 mM EGTA included in some experiments. Signals were amplified (Multiclamp 700B, Molecular Devices), filtered (10 kHz Bessel), and digitized (16-bit resolution; Digidata 1322A, Molecular Devices).

Passive membrane properties – input resistance (*R*_i_) and cell capacitance (*C*_m_) – were measured in whole-cell voltage clamp (holding potential: −70 mV; −5 or −10 mV pulses at 200 Hz or −5 mV pulses lasting 20 ms). Cells were excluded if pipette access resistance (*R*a) exceeded 20 MΩ or if action potentials failed to overshoot by ≥10 mV.

Active properties were assessed in current clamp using 435 ms depolarizing current pulses (10 pA increments, up to 60 pulses or until action potential degradation occurred, marked by progressive widening and amplitude reduction). Cells were held at −80 to −87 mV, as hyperpolarization was required to resolve low-threshold depolarizations distinguishing V1^FoxP2^ interneurons from V1^R^ or V1^MafB/Pou6f2^ subtypes.

Firing phenotypes were classified as bursting or non-bursting, with spike frequency adaptation (SFA) quantified via instantaneous frequency (IF) plots (peak-to-peak interspike intervals). Bursting cells exhibited high initial IFs that rapidly declined to steady state (horizontal lines: no SFA; downward slopes: SFA). Threshold was defined as the membrane voltage at peak depolarization acceleration during the first suprathreshold pulse. Low-threshold depolarizations preceding spiking were noted where present.

### Single-cell model calibration

We developed a biophysically detailed multicompartment model of the rodent V1^FoxP2^ interneuron based on a previously developed interneuron model^88^ consisting of 5 compartments (soma, basal dendrite and three apical dendrites), and the following ion channel currents: fast Na^+^ current with a slow inactivation gate, delayed-rectifier K^+^ current, fast Ca^2+^-activated K^+^ current and N-type Ca^2+^ current in the soma; and fast Na^+^ and slow K^+^ currents in the dendrites.

Them odel was implemented using the NEURON simulation engine^89^ (https://www.neuron.yale.edu/neuron/) and the NetPyNE multiscale modeling tool^90^ (https://netpyne.org). Model parameters (channel conductance, compartment axial resistance, leak conductance, reversal potential, membrane capacitance and channel time constants) were calibrated by simulating current injections to reproduce the average frequency-current (f-I) curve of the experimental *in vitro* data. For this process, two automated optimization algorithms included in NetPyNE were employed: an evolutionary algorithm and the Tree-structured Parzen Estimator (TPE) Bayesian algorithm^91^. Success of the optimization algorithm was defined by the following criteria: (1) an average absolute difference between experimental and model f-I curves smaller than 1 Hz, (2) biophysical parameters including capacitance, input resistance, AP threshold and resting potential, within the physiological range (see Table1), and (3) physiologically realistic voltage traces, assessed through visual inspection of the plateau potential and AP amplitude.

The model repository is available at: https://github.com/RomanB22/January2025_VIPBased_FoxP2

### Simulation of corticospinal input

To simulate corticospinal input to the V1 interneuron model, we first created a pool of all trial data from CSNs displaying increased activity during lever pushes, and another of all trial data from CSNs displaying decreased activity (4 s peri-movement window, Hit trials only). Their inferred instantaneous spike rate was converted to a list of simulated spike times generated by an inhomogeneous Poisson process. This result was validated by comparing the number of spikes obtained from our generator with the expected number of spikes from the CASCADE algorithm (125 ± 183 spikes per trial for our Poisson generator versus 129 ± 190 estimated by CASCADE). Since the CASCADE algorithm infers spike rates from calcium transients, it does not account for spikes that set the resting fluorescence of each cell. Therefore, an additional resting spike rate of 7 Hz was added, matching the median firing rate of L5B pyramidal neurons during quiet wakefulness^92^. Calcium imaging data suggested reduced activity consists of both a reduction in the frequency and amplitude of spontaneous calcium transients, in addition to a reduction in resting fluorescence. When considering changes in spontaneous calcium transients, we observed a linear relationship between the change in ΔF/F_0_ and the associated change in instantaneous firing rate, and thus could simply estimate the change in firing rate underlying the reduction in baseline fluorescence (median Δ firing rate = −13%). Consequently, for decreased CSNs the resting firing rate was reduced from 7 Hz to 6 Hz during the movement period.

**Table 1.** Model and experimental comparisons for V1 interneurons.

|  | Model | Experiment<br>(Average ± Stdev) |
| --- | --- | --- |
| Rheobase (pA) | 83 | 81 ± 57 |
| Capacitance (pF) | 72 | 50 ± 18 |
| Input resistance (Mohm) | 276 | 269 ± 102 |
| AP threshold (mV) | -45 | -40 ± 1 |
| Resting potential (mV) | -56 | -48 ± 5 |

To characterize the relationship between CSN input and V1 interneuron output, we first considered CSN activity during a period of 1300 ms prior to cue presentation. Simulations were generated for 1-25 CSN inputs^44^ with each simulation consisting of 500 runs with trials randomly sampled from the pool of all CSNs (increased + decreased).

To simulate movement-related CSN input to V1 interneurons, we considered CSN activity during a 500 ms peri-movement window. Simulations were generated for different combinations of driving (CSN_Inc._) and modulatory (CSN_Spont._ & CSN _Dec._) (1-10 CSNs), each consisting of 500 runs in which trials were randomly sampled from their respective datasets. The gain of the input-output transformation was measured as the slope of the best fit line of the monotonically increasing portion of the relationship between driving input and V1 output (F _out_ = *f*(F _driver_)). The offset was calculated as the difference in driving input required to produce an output firing rate of 20 Hz across the two conditions. The output range was defined as the maximum output firing rate minus the output firing rate when the driver input rate = 0.

## Supplementary Figures

**Figure S1.**
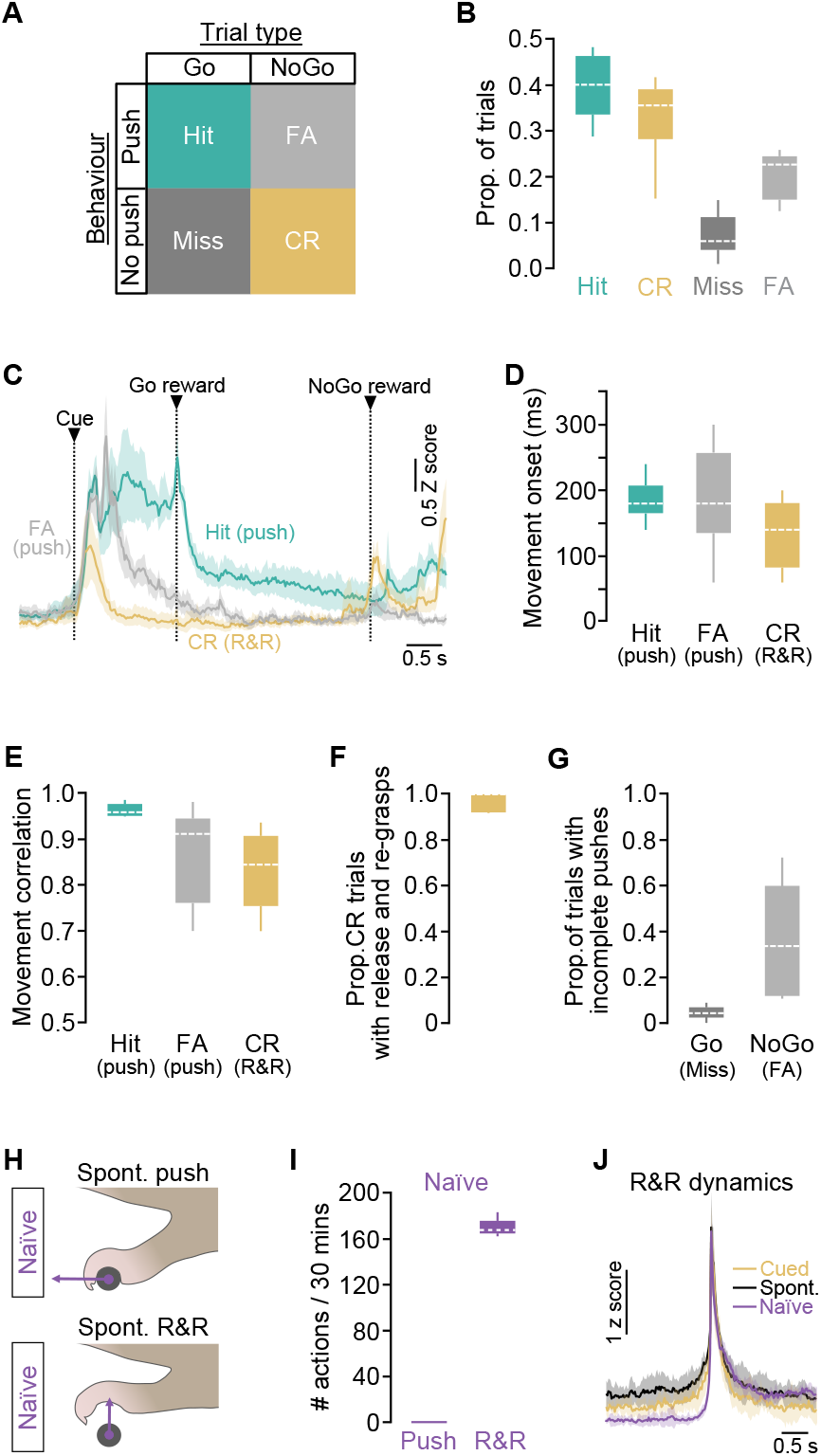
Lever pushes require motor skill learning while release and re-grasps are pre-learned movements. (A)Trial type behavioral outcomes. Hit (push during a Go trial); CR, Correct Rejection (no push during a NoGo trial); FA, False Alarm (push during a NoGo trial); Miss (no push during a Go trial). (B)Proportion of Hit (green), Correct Rejection (CR, yellow), Miss (dark gray) and False Alarm (FA, light gray) trials in expert mice (N = 7 mice). Dashed line, median. (C)Average magnitude and timing of forelimb movements during Hit (push, green), False Alarm (push, light gray) and Correct Rejection (release and regrasp, yellow) trials aligned to cue. Timing of median Go and NoGo reward delivery is shown for comparison. Data represent mean Z scored motion index ± 95% CI (N = 7 mice). (D)Median movement onset time during Hit, False Alarm and Correct Rejection trials (N = 7 mice). Dashed line, median. (E)Average across-trial correlation (pairwise trial-to-trial correlation) of forelimb movements (motion index) during Hit (push, green), False Alarm (FA, push, light gray) and Correct Rejection (CR, release and re-grasp (R&R), yellow) trials (N = 7 mice). Dashed line, median. (F)Proportion of NoGo Correct Rejection trials with release and re-grasp movements (N = 7 mice). Dashed line, median. (G)Proportion of incorrect Go and NoGo trials with incomplete pushes (classified as Go Miss and NoGo False Alarm, respectively) (N = 7 mice). Dashed line, median. (H)Schematic depiction of paw movement trajectories during spontaneous pushes (upper) and release and re-grasps (R&Rs, lower) in naïve mice in the absence of a cue. (I)Number of pushes or release and re-grasps (R&Rs) during a 30 minute behavioral session in naïve mice in the absence of a cue (N = 4 mice). Dashed line, median. (J)Average timing and magnitude of release and re-grasp movements in: expert mice during cued NoGo trials (Cued, yellow); expert mice during the inter-trial interval (i.e. Spont., black); or in naïve untrained mice (naïve, purple). Data represents mean z-scored motion index ± 95% CI (N = 7, 7 and 4 mice, respectively).

**Figure S2.**
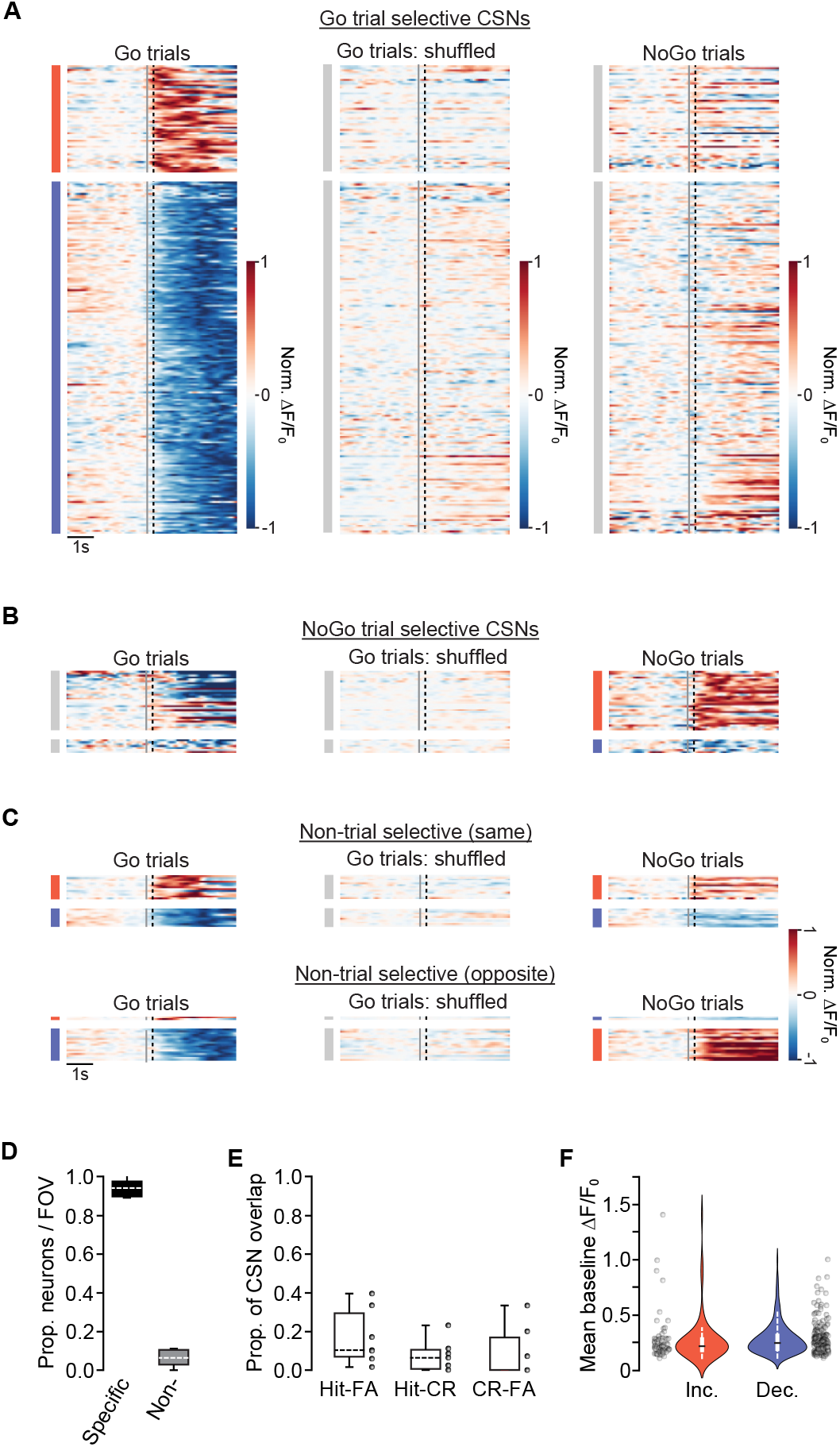
Trial type-specific CSN activity patterns. (A)Normalized average ΔF/F_0_ in Go trial selective CSNs during Go (push) and NoGo (release and re-grasp) trials aligned to movement initiation (black dashed line). Note, Go trial-selective CSNs constitute a subpopulation of the CSNs shown in Figure 1E. Each row represents the average activity of an individual CSN compared across trial types. Groupings: increased activity during Go trials (red bar, n = 56/239 neurons); decreased activity during Go trials (blue bar, n = 183/239); shuffled Go trial data and corresponding activity during NoGo trials (gray bars, 239/239 neurons). In total, n = 239 neurons from 7 fields of view, N = 7 mice. Gray solid line, median cue presentation. (B)Same as shown in (A) but for NoGo trial selective CSNs. Groupings: increased activity during NoGo trials (red bar, n = 31/38 neurons); decreased activity during NoGo trials (blue bar, n = 7/38); corresponding activity during Go trials (gray bars, 38/38 neurons). In total, n = 38 neurons from 7 fields of view, N = 7 mice. Gray solid line, median cue presentation. (C) Same as shown in (A) but for non-trial selective CSNs (*Top*, same direction of change; *Bottom*, opposite direction of change). Groupings: increased activity during both trial types (red bars, n = 13/23 neurons); decreased activity during both trial types (blue bars, n = 10/23); increased activity during Go trials and decreased activity during NoGo trials (n = 2/19 neurons); decreased activity during Go trials and increased activity during NoGo trials (n = 17/19). In total, n = 42 neurons from 7 fields of view, N = 7 mice. Gray line, median cue presentation. (C)Proportion of CSNs displaying movement specificity (N = 7 mice). Specific, movement specific; Non-, non-movement specific. Dashed line, median. (D)Proportion of CSNs that show activity across different movement types (Hits & FAs, Hits & CRs, CRs & FAs) (N = 7 mice). Black circles, individual mice. Dashed line, median. (E)Distribution of mean baseline ΔF/F_0_ in CSNs showing increased (red) or decreased (blue) activity upon cue presentation. Note Inc. and Dec. include both Go and NoGo specific CSNs. Black circles represent individual cells. Black bar, median.

**Figure S3.**
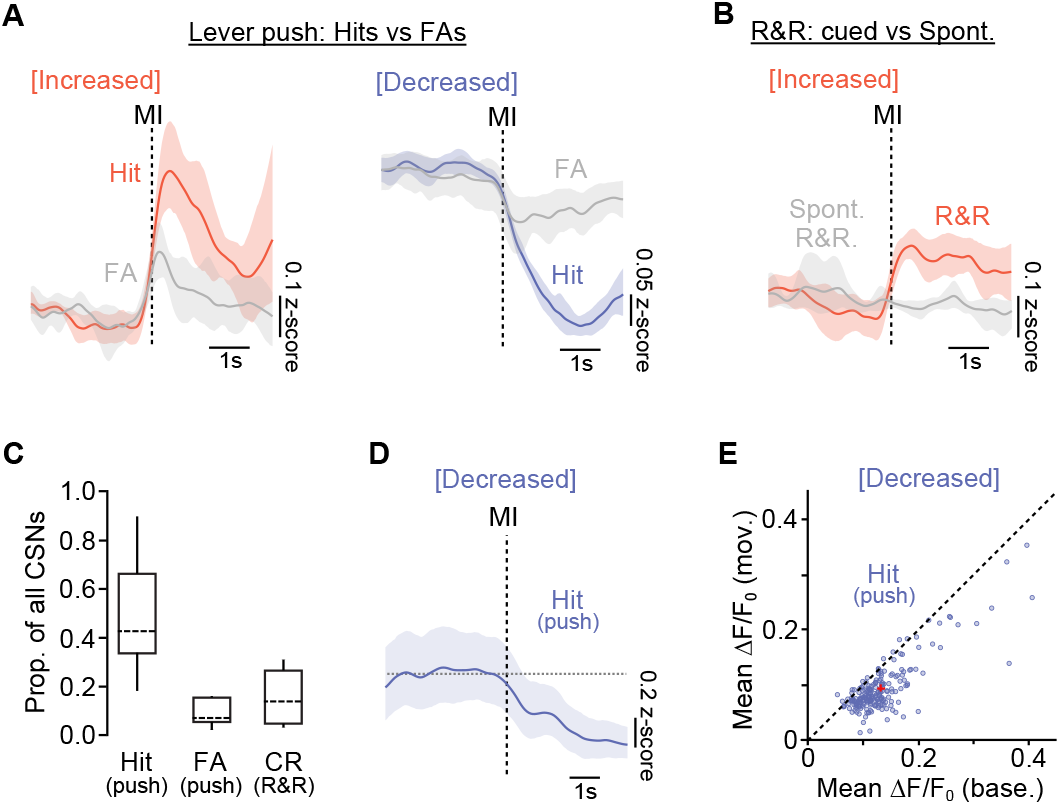
Trial-type specific CSN population responses. (A)Z scored population ΔF/F_0_ ± 95% CI during Hit (lever push after Go cue, red or blue) and False Alarm (lever push after NoGo cue, gray) trials in CSNs that show increased (*left*, red) or decreased (*right*, blue) activity aligned to movement initiation (MI, black dashed line). (B)Z scored population ΔF/F_0_ ± 95% CI during Correct Rejection (release and re-grasp, red) trials and spontaneous release and re-grasps (no cue, gray). MI, movement initiation (black dashed line). (C)Proportion of all CSNs active during Hit (push), False Alarm (push) and Correct Rejection (release and re-grasp) trials in expert mice (N = 7 mice). Dashed line, median. (D)Z scored population ΔF/F_0_ ± 95% CI during Hit push trials (blue) in the absence of bursts of spontaneous events in baseline. MI, movement initiation (black dashed line). (E)Z scored ΔF/F_0_ ± 95% CI during baseline (base.) and movement (mov.) in CSNs that decreased their activity during Go push trials (n = 210 neurons, N = 7 mice). Red cross, median. Dashed line, unity.

**Figure S4.**
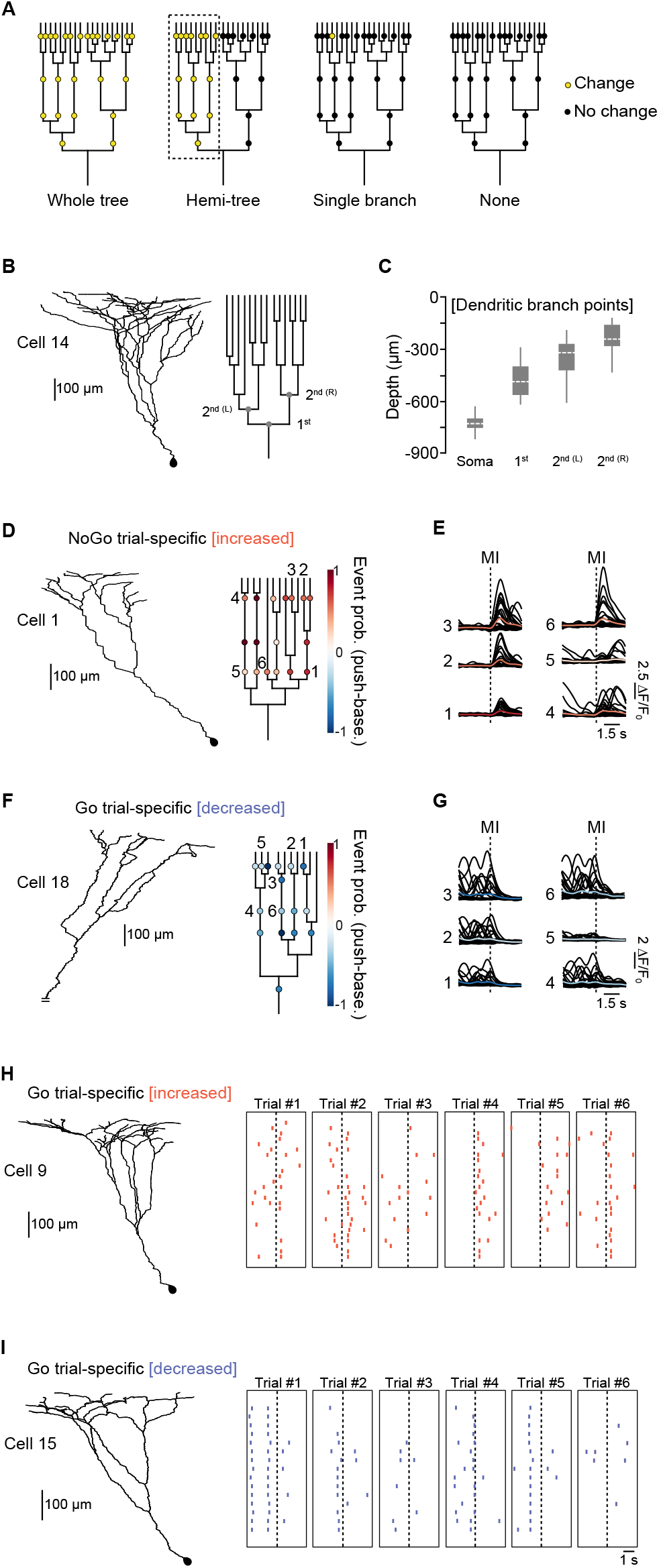
C6-projecting CSN morphometry and example trial-type-specific dendritic activity patterns during Go and NoGo trials. (A)Dendrograms depicting activity patterns classified as whole tree, hemi-tree (purple dashed box), single branch and none. (B)*Left*, reconstruction of the trunk and apical dendrites of a representative CSN. *Right*, corresponding dendrogram showing depth of the first and second (left and right) dendritic branch points. (C)Box plots showing the depth of C6-projecting CSN somata and first and second (left and right) dendritic branch points (n = 16 neurons, N = 9 mice). Dashed line, median. (D)*Left*, reconstruction of the trunk and apical dendrites of a NoGo trial-specific CSN displaying increased activity during release and re-grasps. *Right*, corresponding dendrogram showing location of imaged dendrites and change in event probability across trials (i.e. event probability during push - event probability during baseline period). (E)Overlaid raw ΔF/F_0_ traces (black) and mean (red) from individual dendrites numbered in panel (D) (32 consecutive trials). (F)*Left*, reconstruction of the trunk and apical dendrites of a Go trial-specific CSN displaying decreased activity during lever pushes. *Right*, corresponding dendrogram showing location of imaged dendrites and change in event probability across trials (i.e. event probability during push - event probability during baseline period). (G)Overlaid raw ΔF/F_0_ traces (black) and mean (red) from individual dendrites numbered in panel (F) (31 consecutive trials). (H)*Left*, reconstruction of the trunk and apical dendrites of a Go trial-specific CSN displaying increased activity during lever pushes. *Right*, raster showing spontaneous and cue-evoked calcium transients across trials. Each row represents an individual dendritic branch; each point on the raster an individual calcium event. Dashed line, movement initiation. (I)*Left*, reconstruction of the trunk and apical dendrites of a Go trial-specific CSN displaying decreased activity during lever pushes. *Right*, raster showing spontaneous and cue-evoked calcium transients across trials. Each row represents an individual dendritic branch; each point on the raster an individual calcium event. Dashed line, movement initiation. (J)*Left*, reconstruction of the trunk and apical dendrites of a Go trial-specific CSN displaying decreased activity during lever pushes. *Right*, corresponding dendrogram showing location of imaged dendrites and change in event probability across trials (i.e. event probability during push - event probability during baseline period). (K)Overlaid raw ΔF/F_0_ traces (black) and mean (red) from individual dendrites numbered in panel (F) (31 consecutive trials). (L)*Left*, reconstruction of the trunk and apical dendrites of a Go trial-specific CSN displaying increased activity during lever pushes. *Right*, raster showing spontaneous and cue-evoked calcium transients across trials. Each row represents an individual dendritic branch; each point on the raster an individual calcium event. Dashed line, movement initiation. (M)*Left*, reconstruction of the trunk and apical dendrites of a Go trial-specific CSN displaying decreased activity during lever pushes. *Right*, raster showing spontaneous and cue-evoked calcium transients across trials. Each row represents an individual dendritic branch; each point on the raster an individual calcium event. Dashed line, movement initiation.

**Figure S5.**
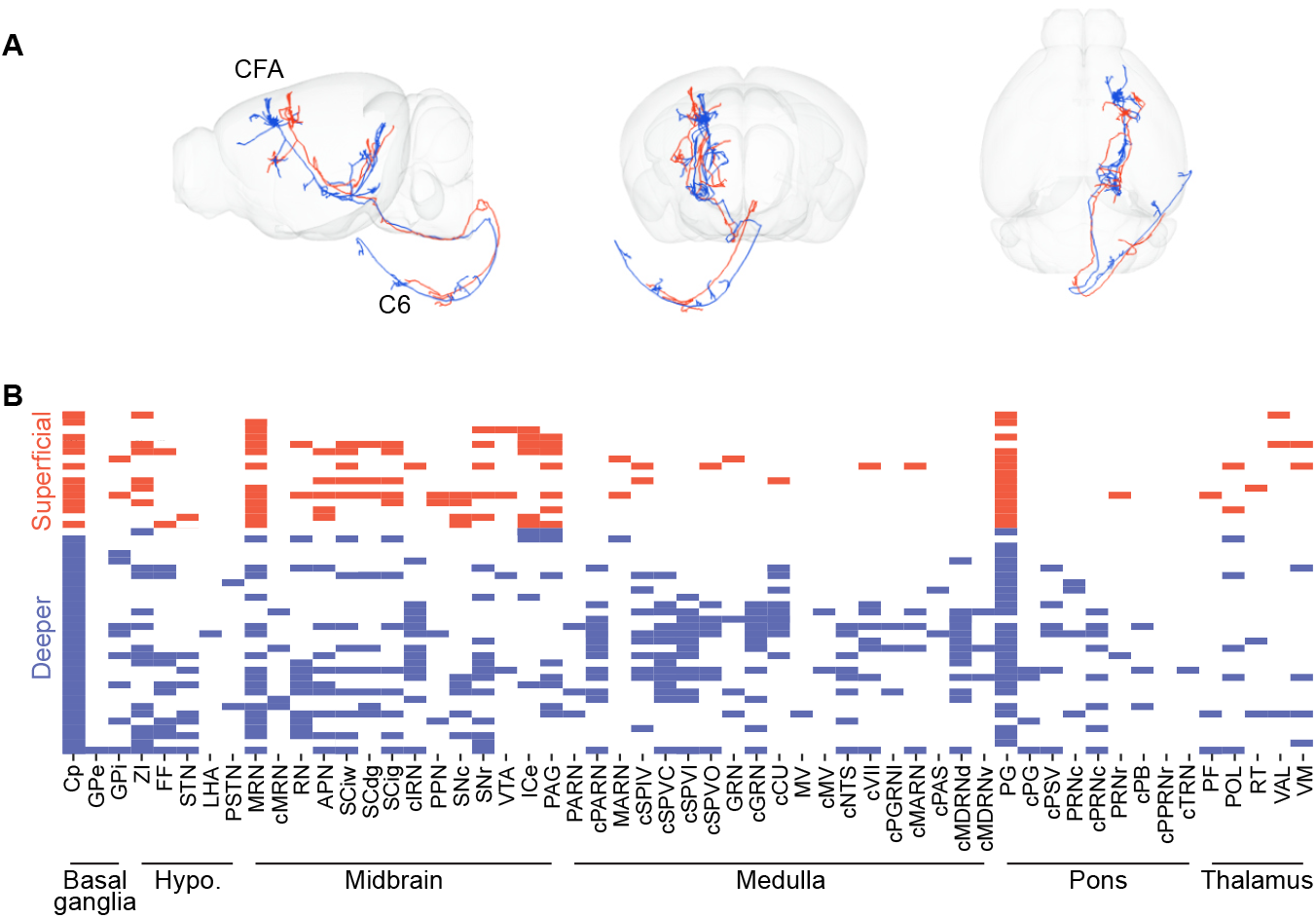
Supraspinal projection targets of caudal forelimb region CSNs. (A)Example visualizations of supraspinal projection targets of two CSNs from superficial (red) and deeper (blue) regions of CFA. CFA, caudal forelimb region of motor cortex; C6, C6 region of the spinal cord. (B)Raster plot showing proportion of CSNs in superficial (red) and deeper (blue) regions of CFA that target different supraspinal brain regions (n = 47 CSNs, N = 4 mice). CSNs are ordered by depth from the pial surface. Note the scarcity of projections from superficial CSN in the medulla.

**Figure S6.**
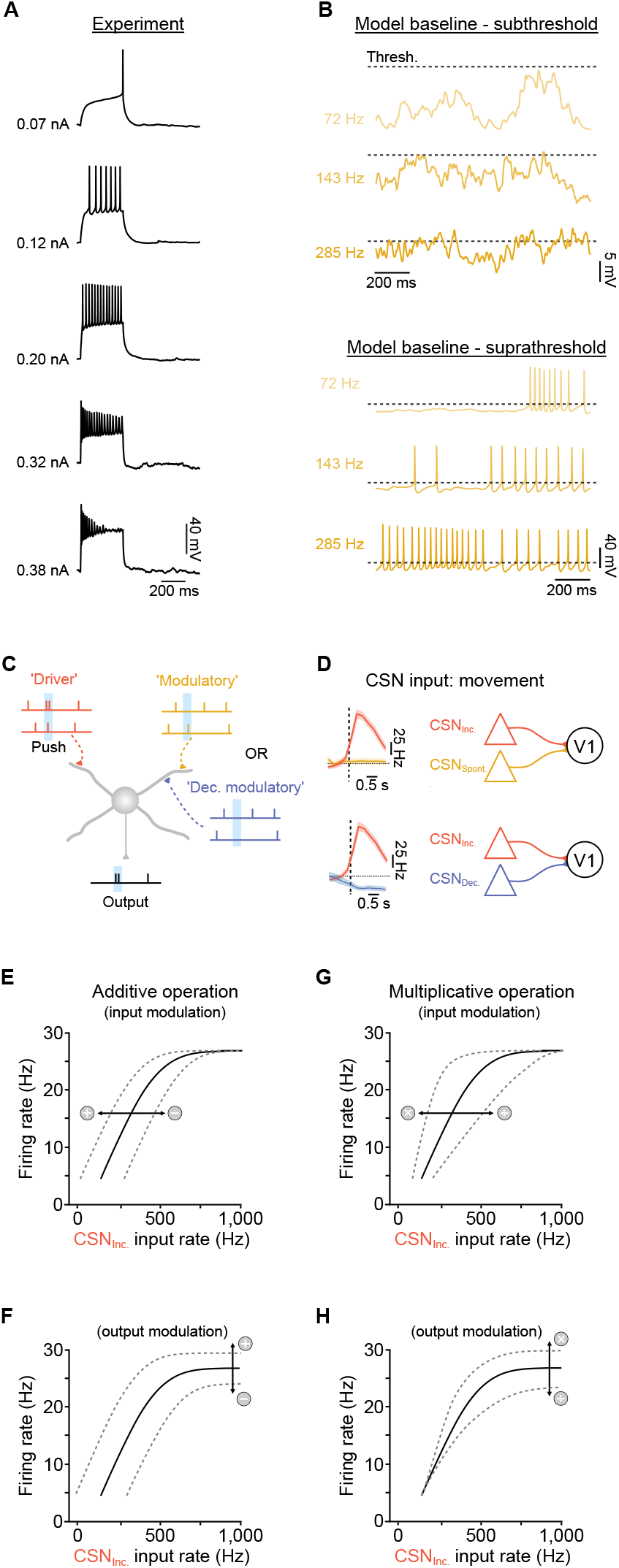
Voltage changes and transformations of input-output relationships in V1 interneurons in vitro and in silico. (A)Example voltage clamp recording from a V1 interneuron *in vitro*. Increasing current injection drives increased spike rate until depolarisation block > 0.38 nA. (B)*Top*, V1 model: baseline membrane potential with respect to threshold (dashed line) when modeling increasing rates of CSN input. Model spiking mechanism has been removed to illustrate subthreshold dynamics. *Bottom*, V1 model: baseline firing rate when modeling increasing rates of CSN input. Dashed line, threshold. (C)Schematic depicting coincident excitatory ‘driver’ input during lever pushes (CSN_Inc._, red) and spontaneous ‘modulatory’ input from layer 5B CSNs that do not change their firing rate during movement (CSN_Spont._, yellow). The effects of changing the modulatory input were modelled as a decrease in spontaneous input during movement (CSN_Dec._, blue) to mimic effects observed *in vivo*. (D)Schematic depicting two V1 model input conditions. *Top*, combining a subpopulation of CSNs that display increased activity during movement (red, CSN_Inc._) with a subpopulation that maintained baseline, spontaneous firing rate throughout (CSN_Spont._, yellow). *Bottom*, combining a subpopulation of CSNs that display increased activity during movement (red, _CSNInc_.) with a subpopulation that decreased activity during movement (CSN_Dec._, blue), mimicking *in vivo* conditions. (E-F) If both driver and modulatory inputs are linearly summed, changing the modulatory input will shift the input-output relationship along the x-axis without changing the shape (E). This is referred to as an additive operation. Note, the shift can be either positive or negative. An additive operation can also be performed on the output of a neuron which shifts the relationship along the y-axis (F). Schematics adapted from Silver (2010)^47^. (G-H) If driver and modulatory inputs are multiplied together during integration, changing the modulatory input will change the slope or gain of the input-output relationship. This is referred to as a multiplicative operation and can change in the input-output scaling along the (G) x-axis (input modulation) or (H) y-axis (output modulation). Schematics adapted from Silver (2010)^47^.

